# Proximity-labeling reveals novel host and parasite proteins at the *Toxoplasma* parasitophorous vacuole membrane

**DOI:** 10.1101/2021.02.02.429490

**Authors:** Alicja M. Cygan, Pierre M. Jean Beltran, Tess C. Branon, Alice Y. Ting, Steven A. Carr, John C. Boothroyd

**Author notes:** Department of Molecular and Cell Biology, University of California, Berkeley CA, USA. Address correspondence to John C. Boothroyd.

## Abstract

*Toxoplasma gondii* is a ubiquitous, intracellular parasite that envelopes its parasitophorous vacuole with a protein-laden membrane (PVM). The PVM is critical for interactions with the infected host cell such as nutrient transport and immune defense. Only a few parasite and host proteins have so far been identified on the host-cytosolic side of the PVM. We report here the use of human foreskin fibroblasts expressing the proximity-labeling enzyme miniTurbo, fused to a domain that targets it to this face of the PVM, in combination with quantitative proteomics to specifically identify proteins present at this crucial interface. Out of numerous human and parasite proteins with candidate PVM localization, we validate three novel parasite proteins (TGGT1_269950, TGGT1_215360, and TGGT1_217530) and four new host proteins (PDCD6IP/ALIX, PDCD6, CC2D1A, and MOSPD2) as localized to the PVM in infected human cells through immunofluorescence microscopy. These results significantly expand our knowledge of proteins present at the PVM and, given that three of the validated host proteins are components of the ESCRT machinery, they further suggest that novel biology is operating at this crucial host-pathogen interface.

**Importance:** *Toxoplasma* is an intracellular pathogen which resides and replicates inside a membrane-bound vacuole in infected cells. This vacuole is modified by both parasite and host proteins which participate in a variety of host-parasite interactions at this interface, including nutrient exchange, effector transport, and immune modulation. Only a small number of parasite and host proteins present at the vacuolar membrane and exposed to the host cytosol have thus far been identified. Here we report the identification of several novel parasite and host proteins present at the vacuolar membrane using enzyme-catalyzed proximity-labeling, significantly increasing our knowledge of the molecular players present and novel biology occurring at this crucial interface.

## Introduction

*Toxoplasma gondii,* the prototypical model organism for the phylum Apicomplexa, is uniquely capable of infecting virtually any nucleated cell in almost any warm-blooded animal. This ubiquitous intracellular parasite can be especially dangerous for immunocompromised individuals, the developing fetus, and is also a significant veterinary pathogen (1). In addition to its medical importance, *Toxoplasma* is a powerful model organism for studying the biology of intracellular parasitism, due to its genetic tractability and ease of culture.

As an obligate intracellular organism, *Toxoplasma* relies on host cell resources in order to replicate. Like other major pathogenic species of the phylum Apicomplexa (e.g., *Cryptosporidium, Plasmodium), Toxoplasma* has evolved to extensively modify its intracellular environment in order to establish a replicative niche, the parasitophorous vacuole (PV), in its host. The PV is necessary to facilitate *Toxoplasma’s* intracellular development and for the avoidance of host defenses. Among other pathogenic processes, *Toxoplasma* tachyzoites, the rapidly dividing forms of the parasite responsible for acute infection, remodel the host cytoskeleton (e.g., vimentin and microtubule organizing center (2–4)), recruit host organelles (e.g., mitochondria and endoplasmic reticulum (5, 6)), scavenge essential nutrients and host resources (e.g., tryptophan, cholesterol, and iron (7–9)), actively manipulate the host transcriptome to support increased metabolism and inhibit the innate immune response (e.g., accumulation of the host “master regulator” c-Myc (10, 11)), and even physically inactivate host protein defenses mounted at the PV (e.g., inactivation of IFNγ-induced immunity-related GTPases in murine cells (12, 13)).

Much of this host cell manipulation relies on parasite effector proteins (e.g., ROP18, GRA15, GRA17/23, MAF1) introduced into the host during infection (recently reviewed in (14)), and located at the parasitophorous vacuole membrane (PVM), the interface between the developing parasites and host. Exposed to the host cytosol, these parasite effectors can interact with various host processes (recently reviewed in (15)). Despite the critical importance of the parasite-host interactions occurring at the PVM, only a small number of proteins at this boundary have thus far been identified. This lack of knowledge has hampered our ability to mechanistically study interesting and potentially disease-relevant aspects of biology, including processes that might be amenable to parasite- and host-targeted drugs.

To address this lack, we have utilized advances in spatial proteomics to identify host and parasite proteins specifically present at the host cytosolic side of the PVM. Proximity labeling-based methods have successfully been used to identify proteins within various *Toxoplasma* cellular compartments, including the PV lumen (16, 17), relying on expression of the crucial biotin-ligase enzymes fused to parasite-expressed proteins. To date, however, no study has used this approach to identify novel proteins specifically present on the host-cytosolic face of the PVM. By fusing the promiscuous biotin-labeling protein, miniTurbo (18), to a domain that targets it to the PVM and engineering host cells to express this fusion, we have been able to generate high confidence lists of candidate host and parasite proteins present at the host-cytosolic face of the PVM. From a subset of this candidate pool, we have validated three parasite and four host proteins as so localized, including several that indicate previously undescribed processes occurring at this all-important interface.

## Results

### Validation of the miniTurbo system in infected HFFs

To identify host and parasite proteins exposed to the host cytosol at the PVM, we utilized enzyme-catalyzed proximity-labeling via the promiscuous biotin ligase, miniTurbo, a recently developed variant of BioID (18, 19). To target miniTurbo to the PVM, we fused V5-tagged miniTurbo-NES (NES = nuclear export signal) to the arginine-rich-amphipathic helix (RAH) domain (amino acids 104-223) of the *Toxoplasma* ROP17 protein (**Fig. 1A**), which had previously been shown to be capable of localizing mCherry to the PVM when heterologously expressed in human cells (20). We stably introduced this construct, as well as V5-tagged miniTurbo-NES without the RAH domain as a control, into human foreskin fibroblasts (HFFs) using lentiviral transduction to generate cell lines stably expressing one or other of the miniTurbo proteins (hereafter referred to as “HFF miniTurbo-RAH” and “HFF miniTurbo”, respectively).

**Figure 1.**
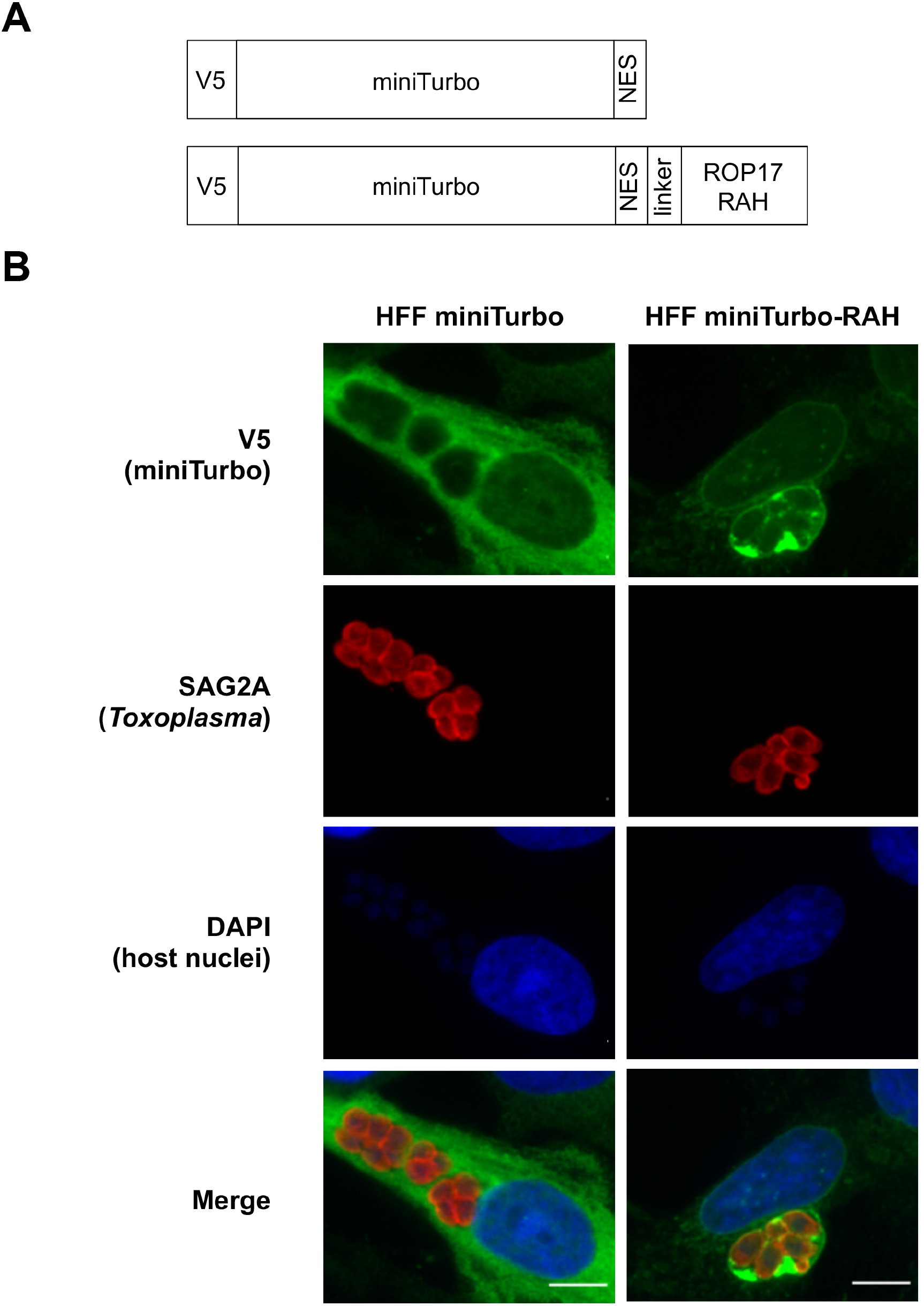
The arginine-rich-amphipathic helix domain (RAH) of ROP17 is sufficient to localize miniTurbo to the *Toxoplasma* PVM in infected cells. **A.** Schematic of the miniTurbo fusion proteins expressed in HFFs depicting the epitope tag (V5), miniTurbo ligase (miniTurbo), nuclear export signal (NES), flexible linker (linker, [amino acids KGSGSTSGSG]), and RAH domain of ROP17 (ROP17 RAH, [amino acids # 104-223]). **B.** Representative immunofluorescence microscopy images of tachyzoite-infected HFF populations stably expressing miniTurbo fusion proteins depicted in panel A. *RHΔhptΔku80* tachyzoites were allowed to infect the corresponding HFFs for 24 hours before the infected monolayers were fixed with methanol. The miniTurbo fusion proteins were detected with mouse anti-V5 antibodies (green), tachyzoites were detected with rabbit anti-SAG2A antibodies, and host nuclei were visualized using DAPI (blue). Scale bar is 10μm.

We assessed the localization of miniTurbo in these cells infected with *Toxoplasma* tachyzoites (RH strain) using immunofluorescence assay (IFA). The results (**Fig. 1B**) show a diffuse, cytosolic staining pattern in HFF miniTurbo cells, consistent with the localization of this construct in other human cell lines (18), and with no apparent association with parasite vacuoles. In HFF miniTurbo-RAH cells, however, miniTurbo staining was highly concentrated at the PVM, being seen even “between” parasites, likely within *Toxoplasma’s* nanotubular intravacuolar network (IVN), the lumens of which are thought to be at least partially contiguous with the host cytosol (20, 21). Thus, the ROP17-RAH domain appears sufficient to target miniTurbo to the PVM in infected cells. In addition, miniTurbo-RAH staining was seen around the nuclear envelope, consistent with the localization previously observed for fluorescent protein fusions with the RAH domains of related *Toxoplasma* proteins (20). This attraction to the host nuclear envelope may be due to the nature in which *Toxoplasma* RAH domains of ROP2 family proteins (i.e., ROP2, ROP5, ROP17, ROP18 etc.) achieve their specificity for the PVM, the exact mechanism of which is unknown but hypothesized to be due to an attraction of the amphipathic helices to the strong negative curvature of the PVM (as a consequence of the IVN) and to a lesser extent attracted to the membranous invaginations of the host nucleus (22). It is notable, however, that the RAH domain is still sufficient to bring fluorescent protein fusions to the PVM in the absence of a mature IVN, leaving open the possibility that a parasite-specific protein or lipid on the PVM may at least partially be driving this association (20).

To test the functionality of the miniTurbo and miniTurbo-RAH fusions in the host cytosol and at the PVM, we infected HFF, HFF miniTurbo, and HFF miniTurbo-RAH cells with tachyzoites, initiated labeling with the addition of exogenous biotin for 1 hour, and assessed biotinylation by western blot and IFA. Streptavidin blot analysis of whole cell lysates (**Fig. 2A**) shows that in HFFs that lack miniTurbo, only a few host and parasite proteins are readily observed, presumably endogenously biotinylated since their detection is not dependent on the addition of exogenous biotin. In contrast, substantial biotin labeling was observed in both miniTurbo- and miniTurbo-RAH cells and this additional signal was dependent on the presence of biotin. Staining for the V5 tag confirmed the presence of miniTurbo of the expected mass in these cells (**Fig. 2A**).

**Figure 2.**
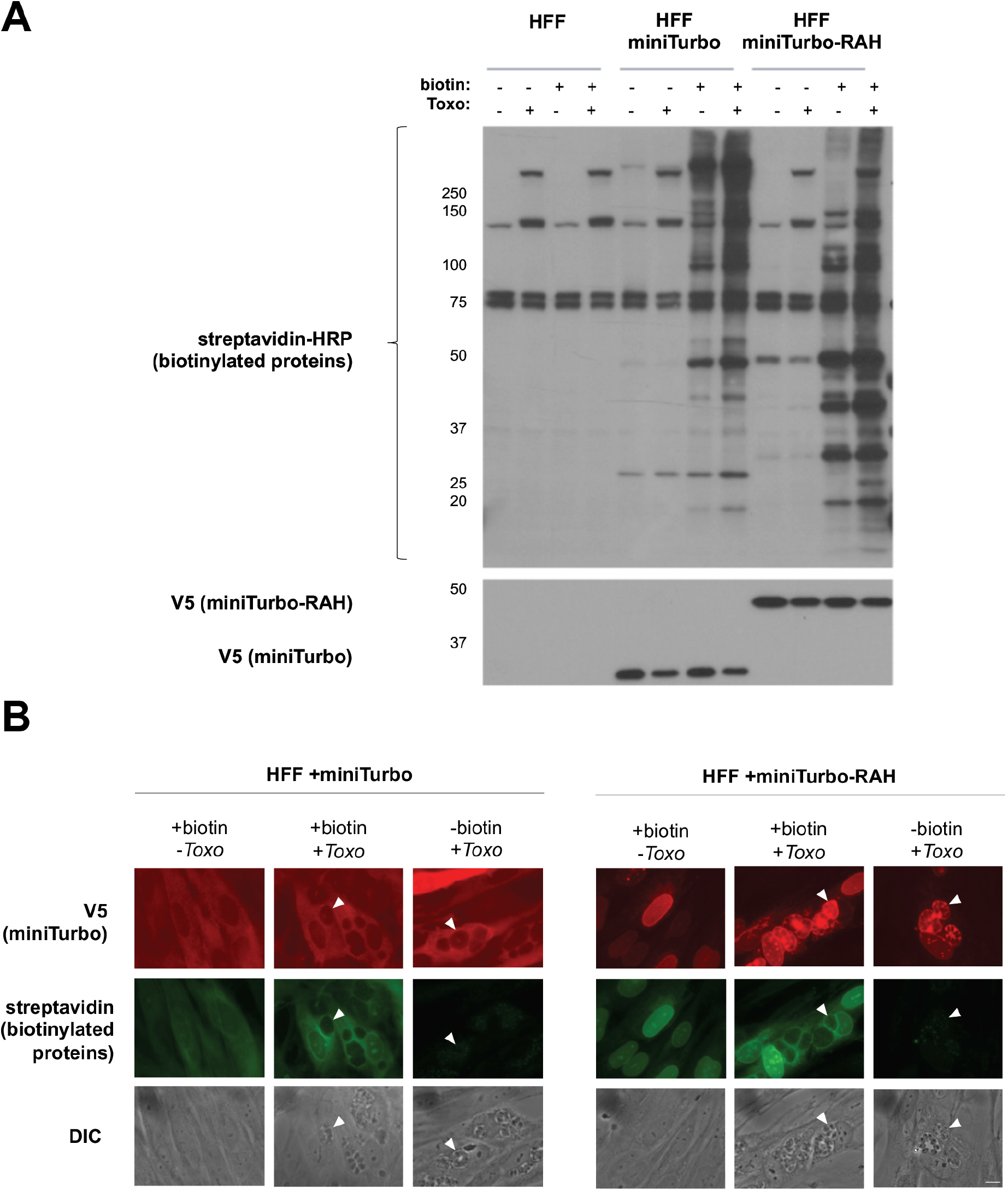
miniTurbo can biotinylate proteins in human foreskin fibroblasts. **A.** Western blot of whole cell lysates. The indicated line of HFFs were either infected or left uninfected (+/−) with *RHΔhptΔku80* tachyzoites for 24 hours and then incubated with or without (+/−) 50 μg of biotin for 1 hour. Biotin labeling was terminated by cooling the cells to 4°C and washing away excess biotin. Whole cell lysates (2 μg) were subsequently prepared and blotted with streptavidin-HRP to visualize biotinylated proteins and mouse anti-V5 antibodies to visualize expression of the miniTurbo fusion proteins. Bands present in the non-miniTurbo expressing HFFs and the HFFs not incubated with biotin represent endogenously biotinylated host and parasite proteins. Approximate migration of a ladder of size standards (sizes in kDa) is indicated. **B.** Representative immunofluorescence microscopy images of HFF populations stably expressing miniTurbo fusion proteins. The indicated line of HFFs were either infected with *RHΔhptΔku80* tachyzoites or left uninfected for 24 hours and subjected to a 1 hour incubation with biotin as described in panel A. After washing, the monolayers were fixed with methanol. The miniTurbo fusion proteins were detected with mouse anti-V5 antibodies (red), biotinylated proteins were detected with streptavidin-HRP (green), and the entire monolayer was visualized with differential interference microscopy (DIC). White arrowheads indicate a representative PV. Scale bar is 10 μm.

To confirm the biotinylating activity in these cells, fluorescent imaging of biotinylated proteins was performed (**Fig. 2B**). This, too, revealed clearly detectable biotinylation in miniTurbo- and miniTurbo-RAH-expressing cells after 1 hour of biotin-labeling that was dependent on the presence of exogenous biotin and largely co-localized with the miniTurbo fusion proteins. In untreated cells, we only observed punctate staining within the parasites, consistent with previous observations of endogenously biotinylated proteins present in the *Toxoplasma* apicoplast (23). Notably, less streptavidin staining was observed “between” parasites where the miniTurbo signal (anti-V5) was strongest in miniTurbo-RAH cells. We speculate that this may be due to a unique redox, pH, or endogenous nucleophile (free amine/lysine availability) environment within the IVN that may influence the activity of miniTurbo, as has been described in other cellular compartments (18). Alternatively, the IVN may be relatively inaccessible to the biotin which might be preferentially utilized by the miniTurbo-RAH present where the IVN nanotubes open into the host cytosol, or this may simply be an artifact due to poor accessibility of streptavidin in fixed cells. Regardless, streptavidin signal was still strongly observed at the PVM in miniTurbo-RAH cells making the system well suited to our goal of exploring this interface.

We next determined the labeling specificity of known PVM proteins by miniTurbo-RAH. We infected HFF miniTurbo-RAH and HFF miniTurbo cells with RHΔ*rop77*::ROP17-3xHA tachyzoites, followed by biotin labeling and enrichment of biotinylated proteins using streptavidin-conjugated beads. Streptavidin blot analysis of the input whole cell lysate, flowthrough from the purification, and eluted proteins showed a strong enrichment for ROP17 within the infected, treated miniTurbo-RAH eluate only, as expected given its known PVM localization (**Fig. 3A**). A much smaller amount of ROP17 was detected in the miniTurbo eluate, consistent with cytosolic miniTurbo having access to ROP17, but to a lesser extent than when concentrated at the PVM (cartoon depiction in **Fig. 3B**). Note that ROP17 exists as both an immature pro-protein and a mature, proteolytically processed protein (24). As expected, the mature form of ROP17 is the one present at the PVM and enriched in the miniTurbo-RAH eluate.

**Figure 3.**
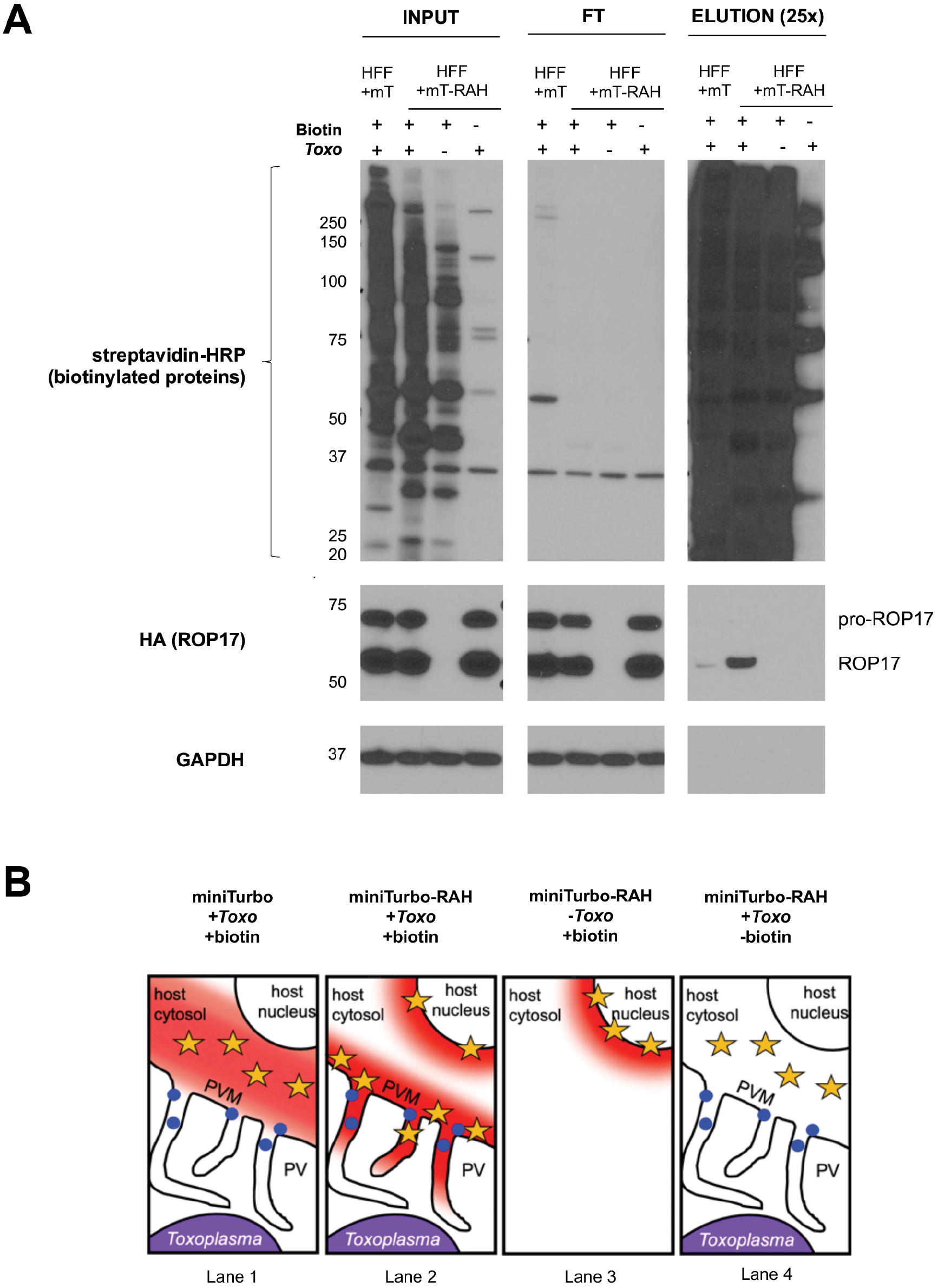
The *Toxoplasma* PVM-localized protein ROP17 is enriched in miniTurbo-RAH cells. **A.** Western blot of whole cell lysates and subsequent elutions after streptavidin affinity purification. The indicated line of HFFs were either infected or left uninfected (+/−) with RH*Δrop77*::ROP17-3xHA tachyzoites for 24 hours and then incubated with or without (+/−) 50 μg of biotin for 1 hour. Whole cell lysates were subsequently prepared and 200 μg of each lysate were incubated with streptavidin-coated magnetic beads overnight. The beads were washed and proteins eluted from the beads by boiling in the presence of 2 mM biotin and BME. Samples of the input lysate (INP), unbound lysate (FT), and eluates (ELU) were separated on an SDS-PAGE gel and blotted with streptavidin-HRP to visualize all biotinylated proteins, mouse anti-HA-HRP antibody to visualize ROP17, and mouse anti-GAPDH to visualize host protein as a loading control. Approximate migration of a ladder of size standards (sizes in kDa) is indicated. **B.** Cartoon depiction of expected regions of biotinylation (depicted by red cloud) near the *Toxoplasma* PVM and their proximity to the localization of ROP17 (blue dot) at the PVM based on miniTurbo fusion localization (yellow star) and addition of biotin.

To confirm and extend these results we next asked whether another PVM-localized parasite protein, MAF1b, is also enriched in the infected miniTurbo-RAH eluate over the miniTurbo eluate. The results (**Fig. S1**) showed this prediction was borne out. Furthermore, a silver stain of the total protein material from the eluates revealed minimal protein background in the miniTurbo-minus and biotin-minus negative controls, indicating that our wash conditions were sufficiently stringent to remove most non-specific background binders (**Fig. S1**). All combined, these results indicate that ROP17-3xHA and endogenous MAF1b, and likely other PVM-associated proteins, are preferentially biotinylated by HFF miniTurbo-RAH vs. HFF miniTurbo cells, and that we could thus use our streptavidin enrichment pipeline followed by quantitative ratiometric analysis to compare protein abundances between the conditions to infer PVM-localization.

### Using the miniTurbo-expressing HFFs to identify novel Toxoplasma PVM proteins

To identify PVM-localized proteins using quantitative proteomics, the HFF cell lines were either mock-infected or infected with RHΔrø*p77*::ROP17-3xHA tachyzoites for 22 hours, biotin was then added for an additional hour, and biotinylated proteins were enriched by affinity purification. Biotinylated proteins captured on the streptavidin beads were digested with trypsin, and the peptides from each experimental condition were then labeled with TMT mass tag reagents, combined, and analyzed by LC-MS/MS (**Fig. 4A**). Western blotting and silver staining from a reserved fraction of these samples showed (**Fig. 4B**) that the washes were sufficient to remove most background protein binding, and that the PVM-localized protein ROP17 was enriched in all four infected miniTurbo-RAH replicates over all other controls. Additionally, principal component analysis of the proteomic dataset showed strong clustering of the biological replicates, indicating high reproducibility (**Fig. 4C**). To quantify differences across experimental groups, we calculated log_2_ fold-changes (log_2_FC) and p-values (moderated t-test) by aggregating all replicates. The full dataset is presented in **File S1**.

**Figure 4.**
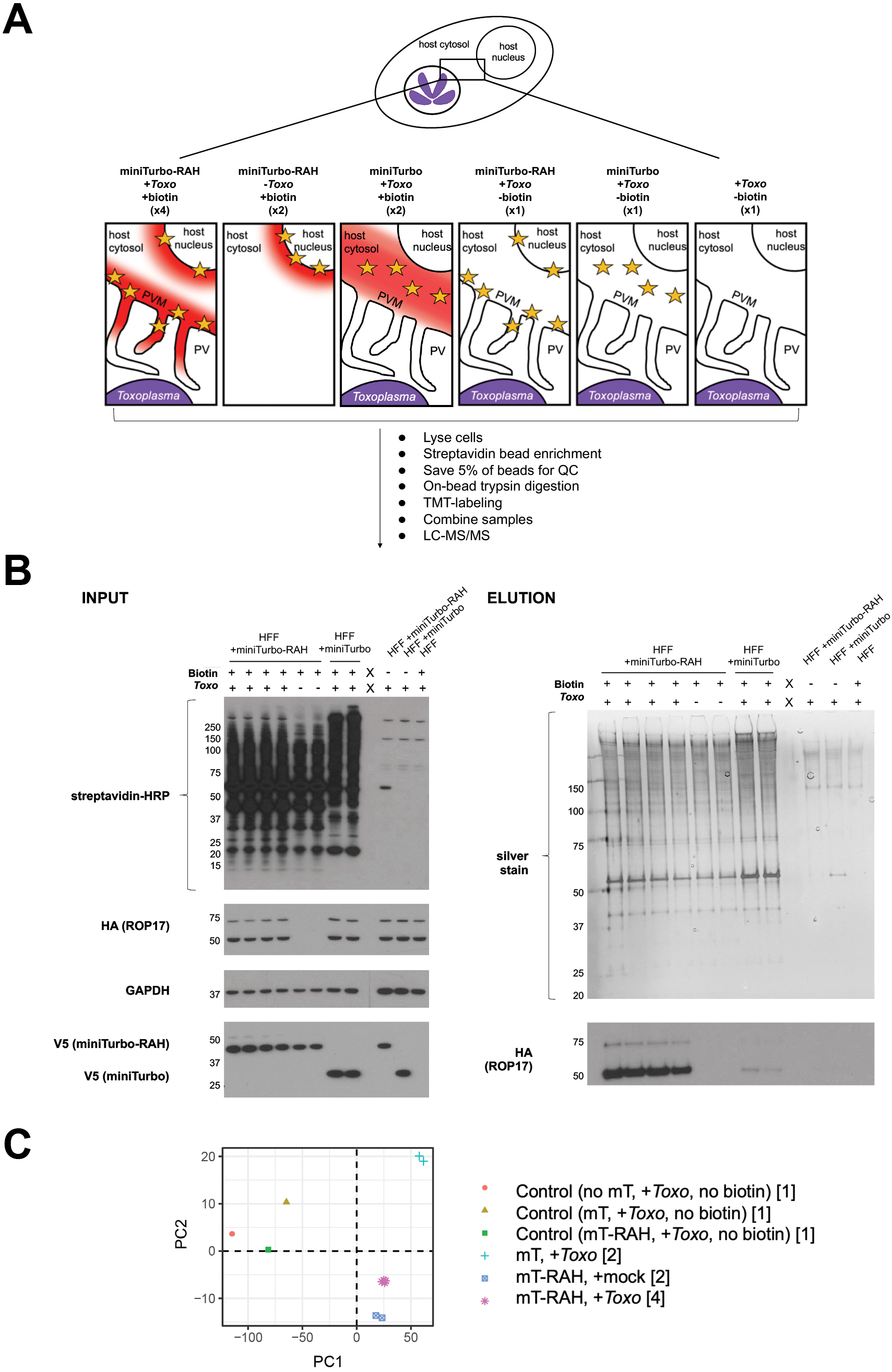
Quantitative mass spectrometry-based proteomic experimental setup and quality control checks. **A.** Overview of proteomics experimental design and cartoon depiction of expected regions of biotinylation (depicted by red cloud) near the *Toxoplasma* PVM based on miniTurbo fusion localization (yellow star) and +/− biotin in the 11 samples submitted for LC/MS-MS. The number of replicates for each sample is indicated in parentheses below the description. The indicated samples were infected with RH*Δrop17*::ROP17-3xHA parasites for 22 hours, and then treated with 50 μM biotin for 1 hour. After treatment and biotin-labeling, cells were lysed and biotinylated proteins enriched with streptavidin magnetic beads, 5% of the beads were saved for a quality control check, the remaining protein was digested on-bead, and subsequent peptides conjugated to TMT labels. All 11 samples were then combined and analyzed by LC-MS/MS. **B.** INPUT: Western blot of whole cell lysates. A sample of each whole cell lysate depicted in panel A was run on a SDS-PAGE gel and blotted with streptavidin-HRP to visualize all biotinylated proteins, mouse anti-HA-HRP antibodies to visualize endogenous ROP17, and mouse anti-V5 antibodies to visualize miniTurbo expression. ELUTION: Silver stain and western blots of eluates from streptavidin affinity purification (from the 5% of beads saved prior to on-bead digestion). The beads were washed and proteins eluted from the beads by boiling in the presence of 2mM biotin and BME. Equivalent volumes of each elution were run on an SDS-PAGE gel and stained with silver to visualize total protein material eluted from the beads. Equivalent volumes of each elution were additionally run on a separate SDS-PAGE gel and blotted with mouse anti-HA-HRP antibodies to check for specific enrichment of endogenous ROP17 in the PVM-miniTurbo localized samples (miniTurbo-RAH +*Toxo*, lanes 1-4). Approximate migration of a ladder of size standards (sizes in kDa) is indicated. **C.** Principal component analysis of the 11 samples analyzed in the proteomics experiment. Numbers in square brackets indicate the number of replicates for each sample type analyzed, as described in panel A.

We refined the candidate PVM-localized proteins by comparison to lists of likely positives (PV/PVM-localized or exported *Toxoplasma* proteins based on published literature) and likely negatives (putatively non-secreted *Toxoplasma* proteins based on a recent LOPIT proteomics dataset (25)). Optimal log_2_FC thresholds were calculated by receiver-operating characteristic (ROC) analysis for the miniTurbo-RAH *+Toxo* / miniTurbo-RAH *+Toxo* (no biotin) comparison (log_2_FC > 2.95) and for the miniTurbo-RAH *+ Toxo* / miniTurbo *+Toxo* comparison (log_2_FC > −0.65) (**Fig. S2**). Using these threshold criteria (**Fig. 5A**, yellow shading), we produced a list of *Toxoplasma* proteins putatively enriched at the PVM (**Table 1**).

**Figure 5.**
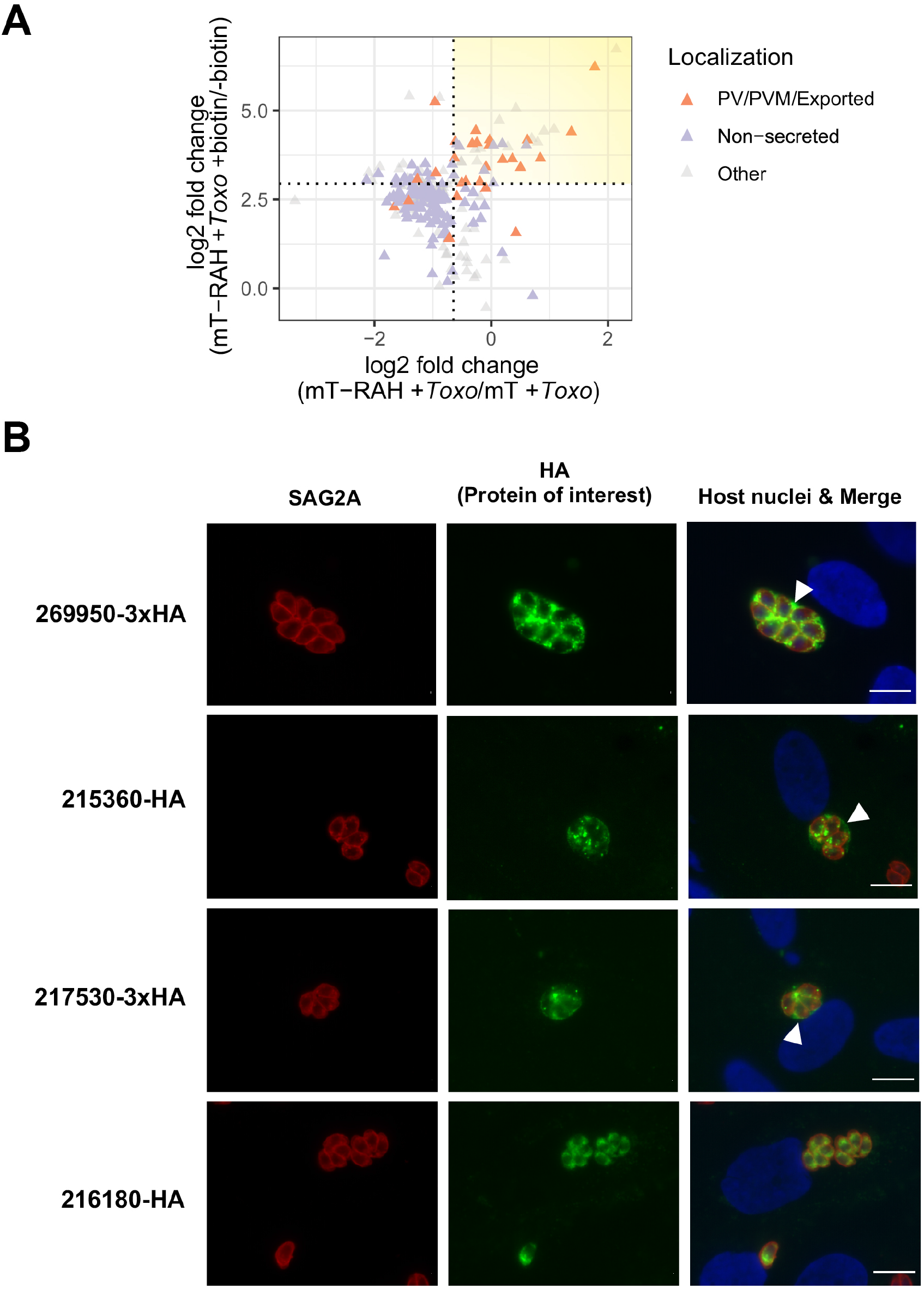
The hypothetical *Toxoplasma* proteins 269950, 215360, and 217530 localize to the *Toxoplasma* parasitophorous vacuole in infected cells. **A.** Two-dimensional plot showing log_2_ fold-changes between miniTurbo-RAH and miniTurbo infected samples (x-axis) and log_2_ fold-changes between +biotin and -biotin miniTurbo-RAH samples (y-axis). *Toxoplasma* proteins known to be exported or PV/PVM-localized or putatively non-secreted (see Materials & Methods) are indicated by orange and purple symbols, respectively. Dotted lines indicate the optimal threshold for separation of known PV/PVM/exported vs. non-secreted *Toxoplasma* proteins. Yellow shading indicates the position on the plot of the proteins of interest described in **Table 1**. **B.** Representative immunofluorescence microscopy images of endogenously tagged (269950-3xHA and 215360-3xHA) or ectopically expressed under native promoter (215360-HA and 216180-HA) parasite proteins. The populations of parasites were allowed to infect HFFs for 20 hours before the infected monolayers were fixed with methanol. The corresponding tagged proteins in parasites that had successfully incorporated the HA-tag were detected with rat anti-HA antibodies, while all tachyzoites were detected with rabbit anti-SAG2A antibodies. Host nuclei were stained with DAPI. Arrowheads indicate vacuoles with HA-staining outside of the parasites. Scale bar is 10 μm.

**Table 1.** List of *Toxoplasma* proteins putatively enriched at the parasitophorous vacuole membrane. The relative abundance of each protein was determined across all samples using the peptide-level TMT quantification and the log_2_ fold change (log_2_ FC) was determined for various sample ratios. The proteins were filtered by a minimum log_2_ FC of 2.95 for the [mT-RAH + *Toxo*]/[mT-RAH + *Toxo* no biotin] ratio and log_2_ FC of −0.65 for the [mT-RAH + *Toxo*]/[mT + *Toxo*] ratio. Displayed here are all the *Toxoplasma* proteins (identified by at least 2 unique peptides, with a protein score >25, and with percent protein coverage >1%) which passed the above thresholds, ranked according to the log_2_ FC between the averaged [mT-RAH + *Toxo*]/[mT-RAH + *Toxo* no biotin] samples, the majority *Toxoplasma* identifiers (TGGT1_; i.e., the proteins that contain at least half of the peptides belonging to a group of proteins that cannot be unambiguously identified by unique peptides), and the descriptive name for each protein (Description). If the protein had been previously localized to the PV/PVM or exported, the reference is indicated. The hypothetical proteins of unknown location and function that were chosen for follow-up localization studies are bolded.

| Gene ID (TGGT1_) | Description | mT-RAH +biotin/<br>mT-RAH -biotin<br>Log<sub>2</sub> FC | PV/PVM Localization |
| --- | --- | --- | --- |
| **269950** | **hypothetical protein** | **6.73** | **PV/PVM (This study)** |
| 220240 | GRA31 | 6.23 | PV/PVM (Nadipuram et al. 2016) |
| 231210 | sarcolumenin/eps15 family protein | 5.06 |  |
| 279350 | hypothetical protein | 4.73 |  |
| **215360** | **hypothetical protein** | **4.48** | **PV/PVM (This study)** |
| 310780 | GRA4 | 4.44 | PV/PVM (Labruyere et al. 1999) |
| **216180** | **hypothetical protein** | **4.42** | **Not secreted (This study)** |
| 309590 | ROP1 | 4.40 | PV/PVM (Saffer et al. 1992) |
| **217530** | **hypothetical protein** | **4.21** | **PV/PVM (This study)** |
| 410370 | MAF1-related | 4.17 |  |
| 258580 | ROP17 | 4.17 | PV/PVM (Ethridge et al. 2014) |
| 210370 | ROP54 | 4.13 | PV/PVM (Kim et al. 2016) |
| 207920 | hypothetical protein | 4.11 |  |
| 229010 | RON4 | 4.10 | PV/PVM (Bradley et al. 2005) |
| 306030 | glutathione s-transferase, n-terminal domain containing protein | 4.07 |  |
| 203600 | GRA50 | 4.07 | PV/PVM (Tu et al. 2020) |
| 249530 | putative exportin 1 | 4.06 |  |
| 289530 | ribosomal protein RPL19 | 4.03 |  |
| 411470 | MAF1-related | 4.02 |  |
| 292130 | ribosomal protein RPL13A | 4.02 |  |
| 215470 | ribosomal protein RPL10A | 3.99 |  |
| 230980 | myosin I | 3.97 |  |
| 311470 | RON5 | 3.94 |  |
| 300100 | RON2 | 3.91 |  |
| 286450 | GRA5 | 3.68 | PV/PVM (Lecordier et al. 1999) |
| 270240 | MAG1 | 3.67 | PV/PVM (Ferguson & Parmley 2002) |
| 254720 | GRA8 | 3.64 | PV/PVM (Carey et al. 2000) |
| 258870B | hypothetical protein | 3.64 |  |
| 411430 | ROP5 | 3.62 | PV/PVM (El Hajj et al. 2006) |
| 306338B | putative dynein gamma chain, flagellar outer arm | 3.58 |  |
| 306060 | RON8 | 3.56 |  |
| 208830 | GRA16 | 3.42 | PV/exported (Bougdoor et al. 2016) |
| 220950 | MAF1b | 3.40 | PV/PVM (Pernas et al. 2014) |
| 314810 | ribosomal protein RPL7 | 3.31 |  |
| 226430 | reticulon protein | 3.19 |  |
| 309120 | ribosomal protein RPL4 | 3.05 |  |
| 215220 | GRA22 | 3.01 | PV/PVM (Okada et al. 2013) |
| 254470 | MYR1 | 3.00 | PV/PVM (Franco et al. 2014) |
| 291930 | RNA recognition motif-containing protein | 2.96 |  |
| 288650 | GRA12 | 2.96 | PV/PVM (Michelin et al. 2009) |

As expected, we observed strong enrichment of several parasite PVM-localized proteins known to be exposed to the host cytosol, including ROP1, MAF1b and MAF1-related proteins, ROP17, ROP54, GRA5, and ROP5 (12, 13, 26–29). Additionally, many proteins (such as GRA31, GRA4, GRA8, and GRA50) which had previously been shown to be secreted into the PV (16, 30–32), and predicted by Phobius (33) to contain transmembrane domains, suggesting integration into the PVM and exposure to the host cytosol, were also enriched. Most importantly, and in addition to these known examples of PVM-localized proteins, several uncharacterized (hypothetical) proteins were also enriched, including TGGT1_269950, TGGT1_215360, TGGT1_216180, and TGGT1_217530, suggesting a likely PVM localization. These four proteins were thus chosen for subsequent analysis of their localization in infected cells. Of note, during the course of this work, TGGT1_269950 was identified as a *Toxoplasma* protein important in conferring parasite fitness in IFNγ-stimulated murine bone marrow-derived macrophages (34), but its intracellular localization in infected cells was not determined in that publication.

To localize these candidate PVM proteins in infected cells, we generated populations of parasites in which these genes were either endogenously modified to encode a 3xHA tag immediately before the stop codon, or in which a C-terminally HA-tagged version of these proteins was ectopically expressed under each gene’s native promoter. Protein localization was then assessed by IFA. The results (**Fig. 5B**) show a clear PV-like signal outside of the parasites in the 269950-3xHA, 215360-HA, and 217530-3xHA populations, including at the periphery of the PV. Thus, we conclude that 269950, 215360, and 217530 are at least transiently localized to the *Toxoplasma* PV/PVM during infection. In contrast, we did not observe any staining outside of the parasites in the 216180-HA population, suggesting that, at least at this time point, this protein is not present in the PVM and it may be a false-positive in our enrichment list.

### Using the miniTurbo-expressing HFFs to identify novel human PVM proteins

In addition to novel parasite proteins, we also aimed to identify host protein candidates at the PVM by focusing on the host proteins enriched upon infection (miniTurbo-RAH *+Toxo/* miniTurbo-RAH +mock) (**Fig. 6A**). First, the list of candidate host proteins was filtered using the log_2_FC threshold established by ROC analysis for the miniTurbo-RAH +*Toxo*/miniTurbo *+Toxo* comparison (log_2_FC >-0.65). Since we could not generate similar lists of likely positive and likely negative host proteins at the PVM due to the lack of knowledge on host proteins at this interface, we instead rank-sorted the host proteins by their log_2_FC upon infection (**Fig. 6A**, yellow shading). The top 50 enriched host proteins are presented in **Table 2**. This list of proteins was analyzed for functional enrichment (**Fig. 6B**) which revealed “viral budding via host ESCRT (endosomal sorting complexes required for transport) complex” to be the most significantly enriched Gene Ontology biological process term. Further analysis looking for known protein-protein interactions (**Fig. 6C**), confirmed an enrichment for several proteins known to physically associate within the ESCRT complex: CC2D1A, PDCD6, PDCD6IP, TSG101, VSP28, and CHMP4B, (**Fig. 6C**) (35).

**Figure 6.**
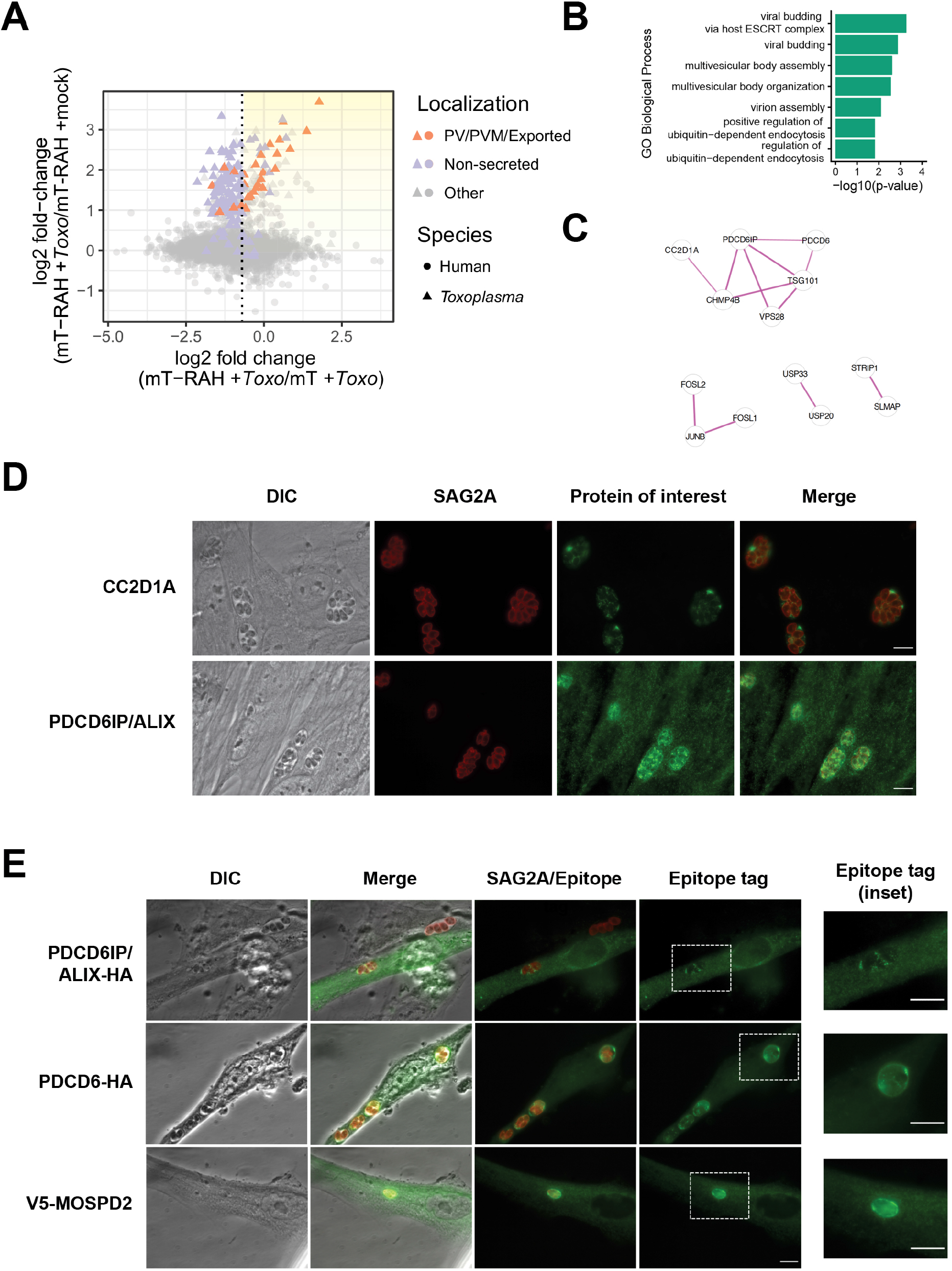
Host ESCRT-associated proteins PDCD6, ALIX, and CC2D1A, and the host ER-organelle tethering protein MOSPD2 localize to the *Toxoplasma* parasitophorous vacuole membrane. **A.** Two-dimensional plot showing log_2_ fold-changes between miniTurbo-RAH and miniTurbo-infected samples (x-axis) and log_2_ fold-changes between infected and mock-infected miniTurbo-RAH samples (y-axis). *Toxoplasma* proteins known to be exported or PV/PVM-localized or putatively non-secreted are labeled using the same color scheme as in Fig. 5A. Dotted line indicates the optimal threshold for separation of known PV/PVM/exported-localized proteins and non-secreted *Toxoplasma* proteins. Yellow shading indicates the position on the plot of the proteins of interest described in **Table 2**. **B.** Significant terms from functional enrichment analysis performed on the candidate host PVM proteins from Table 2. Terms are shown for the Gene Ontology biological process functional database. **C.** Result of STRING v11 (51) protein-protein interaction analysis of the candidate host PVM proteins from Table 2 with singletons (proteins with no association to another protein) removed. The thickness of the lines between proteins indicates the degree of confidence of the interaction. **D.** Representative immunofluorescence microscopy images of the localization of the host proteins CC2D1A and PDCD6IP/ALIX during *Toxoplasma* infection. HFFs were infected with RH parasites for 20 hours before the infected monolayers were fixed with methanol. Rabbit anti-CC2D1A antibodies and rabbit anti-ALIX antibodies were used to detect the corresponding host proteins. Tachyzoites were detected with rabbit anti-SAG2A antibodies and the infected monolayer was visualized with DIC. Scale bar is 10μm. **E.** Representative immunofluorescence microscopy images of the localization of host proteins upon transient overexpression in HFFs. HFFs were transiently transfected with plasmids expressing the indicated tagged proteins and then infected with RH tachyzoites 24 hours post transfection. The parasites were allowed to infect for 16 hours before the monolayers were fixed with methanol. Mouse anti-V5 and rat anti-HA antibodies were used to detect the corresponding host proteins. Tachyzoites were detected with rabbit anti-SAG2A antibodies and the infected monolayer was visualized with DIC. Dashed white boxes indicate the PVs expanded in the insets. Scale bar is 10 μm.

**Table 2.**
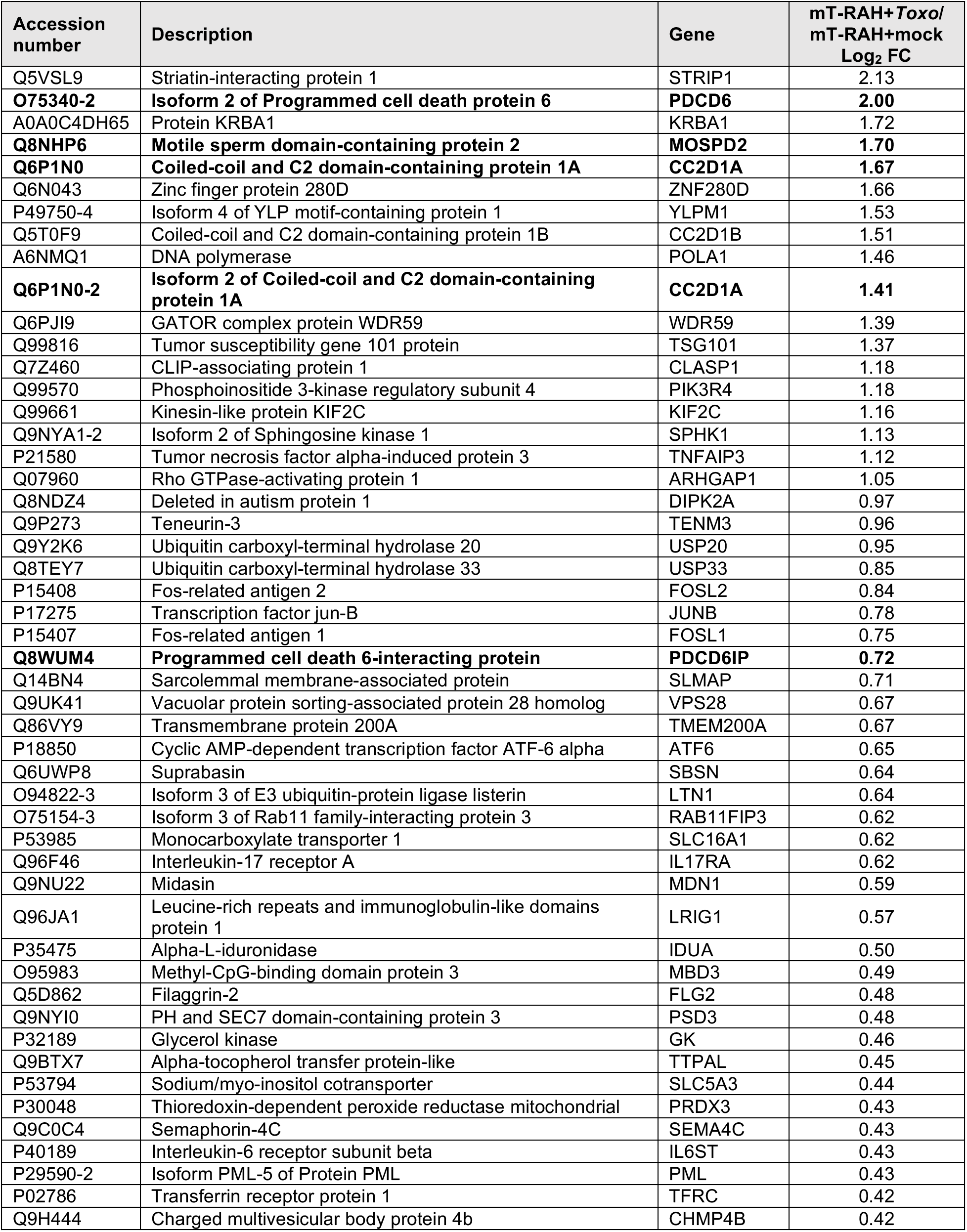
List of the top 50 human proteins putatively enriched at the parasitophorous vacuole membrane. The relative abundance of each protein was determined across all samples using the peptide-level TMT quantification and the log_2_ fold change (log_2_ FC) was determined for various sample ratios. The proteins were first filtered by requiring a minimum log_2_ FC of - 0.65 for the [mT-RAH +*Toxo*]/[mT +*Toxo*] ratio and log_2_ FC of 0 for the [mT-RAH +*Toxo*/mT-RAH *+Toxo,* no biotin] ratio (see materials and methods). Displayed here are the top 50 human proteins (identified by at least 2 unique peptides, with a protein score > 25, and with percent protein coverage >1%) ranked according to the log_2_ FC between the averaged mT-RAH *+ Toxo* samples vs. mT-RAH +mock samples. Shown also are their majority human UniProt identifiers (Accession number), i.e., the proteins that contain at least half of the peptides belonging to a group of proteins that cannot be unambiguously identified by unique peptides, the descriptive name for each protein (Description), and the gene symbol (Gene). The proteins of unknown location in infected cells chosen for follow-up localization studies are bolded.

Host PDCD6IP (also known as ALIX) has previously been reported to associate with at least the nascent PV during the time of *Toxoplasma* invasion (36), as well as the only host protein enriched in a BioID experiment looking to identify proteins within the *Toxoplasma* PV (16). For these reasons, we wanted to confirm if this and the other ESCRT-related host proteins were associated with the PVM at later time points in infection. To do this, we first assessed the localization of PDCD6IP/ALIX and CC2D1A during infection with *Toxoplasma* tachyzoites via IFA using antibodies raised against the endogenous human proteins. The results (**Fig. 6D**) showed clear staining of CC2D1A and PDCD6IP/ALIX surrounding the PVs, as well as present in the spaces “between” parasites, likely within the IVN that often fills such regions. To confirm and extend these results, we also assessed the localization of PDCD6IP/ALIX and PDCD6 (also known as ALG-2), a Ca^2+^ binding protein that interacts with several ESCRT accessory proteins (37), in infected host cells transiently overexpressing C-terminally HA-tagged versions of these proteins. The results (**Fig. 6E**, top two panels) showed moderate, but clear, association of PDCD6/ALIX to the IVN and very strong association of PDCD6/ALG-2 to the PVM.

Another compelling PVM-recruited host candidate that was highly enriched was MOSPD2, an ER-anchored membrane protein that has recently been shown to function as a scaffold to mediate interactions with organelle-bound proteins to form contact sites between the ER and endosomes, mitochondria, or Golgi (38). To confirm if MOSPD2 is recruited to the PVM during infection, we assessed its localization in host cells transiently overexpressing an N-terminally V5-tagged version of MOSPD2. The results (**Fig. 6E**, bottom panel) showed very strong association of MOSPD2 to the PVM. As discussed further below, this could be a clue to the mechanism by which host ER associates with the PVM.

## Discussion

The *Toxoplasma* PVM is a critical platform for host-parasite interactions and by using miniTurbo-catalyzed proximity-labeling we identified candidate host and parasite proteins localized at the host cytosolic side of this interface, and validated several (TGGT1_269950, TGGT1_215360, TGGT1_217530, PDCD6IP/ALIX, PDCD6, CC2D1A, and MOSPD2) as indeed PVM-localized.

A very exciting aspect of this dataset is the new hypotheses that it creates regarding novel host-parasite interactions at the PVM. We confirmed that several host ESCRT accessory proteins are recruited to the PVM during infection, but this leaves unanswered the question of the biological significance of this association. Future work determining whether these host proteins are functioning in a pro- or anti-*Toxoplasma* manner will be critical to understanding how important this recruitment is to parasite development (e.g., how does knockout/knockdown of these components affect parasite replication?). ESCRT proteins mediate many functions within the cell related to membrane remodeling (35, 39), many of which could be relevant to *Toxoplasma.* For example, ESCRT-III (one of the ESCRT subcomplexes) can deform membranes, assemble into long filaments on negatively curved membranes, and promote vesicle budding (35). CC2D1A is a regulator of the ESCRT-III protein CHMP4B (40), and both proteins were highly enriched at the PVM in our dataset. These results suggest the interesting hypothesis that host ESCRT machinery is promoting or maintaining the *Toxoplasma* IVN.

Another interesting hypothesis suggested by our data is that the human protein MOSPD2 may mediate the recruitment of host ER to the PV, given its normal function in bringing the ER in close apposition to other host membranes (38). MOSPD2 interacts with other proteins via “FFAT” motifs [two phenylalanines (FF) in an acidic track (AT)] and analyses assessing the role of *Toxoplasma* proteins possessing such motifs may yield interesting candidates for parasite PVM proteins that may be interacting with MOSPD2. Future immunoprecipitation experiments using infected cells expressing tagged MOSPD2 will be necessary to identify potential parasite binding proteins and further test this hypothesis.

Additionally, future variations of this approach would help to generate even more information as to what proteins are present at the PVM under different conditions. For example, one could assess if there are differences in the proteins present at this interface between different strains of *Toxoplasma,* or even in the closely related organism *Neospora,* and under different stress conditions (e.g., IFNγ treatment). Moreover, combining these data with other recently developed complementary approaches will undoubtedly give us further insight into the molecular composition of the PVM (41).

While the proximity-labeling approach we used is extremely powerful, we’d like to briefly discuss a few important limitations. The first is that the data are likely to be biased towards the most proximal miniTurbo-RAH interactors (i.e., those that are physically closest and spend the most time in the vicinity of the PVM-localized miniTurbo-RAH protein). Given that we used the PVM-localizing RAH domain from ROP17 for this study to bring miniTurbo in proximity to the PVM, it is possible that some of the most enriched *Toxoplasma* proteins identified are direct interactors of ROP17 (a kinase involved in both inactivation of host IFNγ-induced immunity-related GTPases (IRGs) at the PVM as well as protein translocation across the PVM (12, 42)). Indeed, we found ROP5, a ROP17-interactor identified biochemically by tandem affinity purification studies of ROP17 (12), to be enriched within the infected HFF miniTurbo-RAH cells (**Table 1**).

It is interesting then to consider the two most enriched parasite proteins, TGGT1_269950 and GRA31, as possible ROP17-interacting proteins. Future experiments in *Δ269950* and *Δgra31* parasites will be necessary to determine if either are involved in inactivation of host IRGs at the PVM, or translocation of *Toxoplasma* effector proteins across the PVM. It is also possible that one or other of these two proteins could be required for ROP17’s association with the PVM/IVN; i.e., that the ROP17 RAH domain is associating with these proteins which are themselves embedded in the PVM, as an alternative model for how the RAH domain leads to PVM association.

Another important note is that our candidate list of PVM proteins is likely incomplete due to both the limitations of miniTurbo-catalyzed biotinylation and TMT-quantification sensitivity. The biotinyl-5-AMP reactive intermediate that is formed by miniTurbo can only react with primary amines (i.e. free lysines or N-termini). Therefore, if a proximal protein does not have a lysine residue exposed to react with the reactive biotin intermediate, it will not be labeled and we will not detect it. Moreover, even with the use of state-of-the-art MS instrumentation, limitations in the efficiency of biotin enrichment and instrument sensitivity make it possible that we did not observe host cytosol-exposed PVM proteins that are lowly abundant and/or do not have a lysine accessible to the reactive biotin intermediate. Despite these potential limitations, however, this approach has identified many novel molecular players at the PVM.

Altogether, the results reported here demonstrate the power of proximity-labeling coupled with quantitative mass spectrometry to identify proteins exposed to the host cytosol at the *Toxoplasma* PVM, and this rich dataset will enable multiple avenues of future inquiry into crucial host-parasite interactions.

## Materials and Methods

### Parasite strains, culture and infections

All *Toxoplasma* tachyzoites used in this study are in the Type I “RH” background, either *RHΔhptΔku80* (43) or RHΔ*rop17*::ROP17-3xHA (42). These tachyzoites, and all subsequently generated lines, were propagated in human foreskin fibroblasts (HFFs) cultured in complete Dulbecco’s Modified Eagle Medium (cDMEM) supplemented with 10% heat-inactivated fetal bovine serum (FBS; HyClone, Logan, UT), 2 mM L-glutamine, 100 U/ml penicillin and 100 μg/ml streptomycin at 37 °C with 5% CO2. The HFFs were obtained from the neonatal clinic at Stanford University following routine circumcisions that are performed at the request of the parents for cultural, health or other personal medical reasons (i.e., not in any way related to research). These foreskins, which would otherwise be discarded, are fully de-identified and therefore do not constitute “human subjects research”.

Prior to infection, parasites were scraped and syringe-lysed using a 27 G needle, counted using a hemocytometer, and added to HFFs. “Mock” infection was done by first syringe-lysing uninfected HFFs, processing this in the same manner as done for the infected cells, and then adding the same volume of the resulting material as used for infections.

### Plasmid construction

DNA sequences for all fusion constructs used in this study can be found in **File S2**. Briefly, all plasmids were constructed using standard molecular biology techniques. pCDNA-V5-miniTurbo-NES (Addgene plasmid # 107170) (18). The RAH domain of ROP17 was amplified from *Toxoplasma* RH genomic DNA using forward primer 5’-AAGACCCTATTACCGTGATGGGAGGTTGTC-3’ and reverse primer 5’-TCAGCCTATCAAAGGCGGAACTACCGGTG-3’. The pCDNA-V5-miniTurbo-NES-RAH fusion plasmid was generated from this plasmid by overlap extension PCR and a short linker (AAGGGCTCGGGCTCGACCTCGGGCTCGGGA) was introduced between the nuclear export signal (NES) and ROP17 RAH sequence. V5-miniTurbo and V5-miniTurbo-RAH were cloned into the pLenti-CMV-Puro plasmid (a gift from Dr. Jan Carette, Stanford University; Addgene #17452) using Gibson Assembly (NEB).

For *Toxoplasma* endogenous tagging plasmids: ~1500-3000 bp of the 3’-coding sequence of each gene, up to but not including the stop codon, was amplified from RH genomic DNA and cloned into the pTKO2-HPT-3xHA plasmid (44) using Gibson Assembly (NEB).

For *Toxoplasma* ectopic expression plasmids: plasmids to ectopically express HA-tagged proteins off their native promoters were created by PCR amplification of the open reading frame of each gene, minus the stop codon, plus ~2000bp upstream of the start codon to include the native promoter, followed by cloning into the pGRA-HPT-HA plasmid (45).

For human ectopic expression plasmids: Full length cDNA for each gene was amplified from a HeLa cDNA library (Takara Bio) and inserted into a pCDNA expression plasmid modified with either a C-terminal HA or N-terminal V5 tag under the human cytomegalovirus (CMV) promoter using standard molecular biology techniques.

### Toxoplasma transfections

Endogenous tagging plasmids and/or ectopic expression plasmids (with the gene of interest expressed off its native promoter) were transfected into *Toxoplasma* via electroporation using the Amaxa 4D Nuecleofector (Lonza). Tachyzoites were mechanically released in PBS, pelleted, and resuspended in 20 μL P3 Primary Cell Nucleofector Solution (Lonza) with 5-25 μg DNA for transfection. After transfection, parasites were allowed to infect HFFs in DMEM in T25 flasks for 24 hours, after which the medium was changed to complete DMEM supplemented with 50 μg/ml mycophenolic acid and 50 μg/ml xanthine for selection for the hypoxanthine-xanthine-guanine-phosphoribosyltransferae (HXGPRT or HPT) marker for 3-5 days.

### Mammalian cell culture and stable cell line generation

For preparation of lentiviruses, HEK 293T cells in 10cm dishes were transfected at 80% confluency with the lentiviral plasmid pLenti-CMV-Puro (Addgene #17452) containing the gene of interest (2 μg), and the lentiviral packaging plasmids pVSV-G, *pΔVPR,* pAdvant (gifts from Dr. Jan Carette, Stanford University) using FuGENE HD transfection reagent (Promega) according to manufacturer’s instructions. After 24 hours, the medium was replaced with fresh medium. Approximately 48 hours after transfection, the cell medium containing the lentivirus was harvested and filtered through a 0.45-μm filter and supplemented with 8 μg/mL protamine sulfate. To generate stable lines, HFFs in 6-well plates were then infected with the virus-containing medium (1 mL). The following day, the viral-containing medium was removed and replaced with fresh, antibiotic-free medium. The HFFs were allowed to recover for 48 hours and then selected with medium containing 2 μg/mL puromycin for 3 days.

### Mammalian cell transient transfections

HFFs were grown on glass coverslips to ~80% confluency and subsequently transfected with Lipofectamine LTX reagent (Invitrogen) and 500ng of each pCDNA plasmid (with the tagged gene of interest under the human cytomegalovirus promoter) according to the manufacturer’s instructions in antibiotic-free DMEM. Cells were incubated with the transfection reagent for ~16 hours, media changed, and tachyzoites added for another 24 hours.

### Immunofluorescence assay (IFA)

Infected cells grown on glass coverslips were fixed and permeabilized using 100% cold methanol for 10 min. Samples were washed 3x with PBS and blocked using 3% BSA in PBS for 1 hour at room temperature (RT). HA was detected with rat monoclonal anti-HA antibody 3F10 (Roche), SAG2A was detected using rabbit polyclonal anti-SAG2A antibodies (46), V5 was detected with mouse anti-V5 tag monoclonal antibody (Invitrogen R960), human CC2D1A was detected with rabbit anti-CC2D1A antibodies (Sigma HPA005436), human PDCD6IP/ALIX was detected with rabbit anti-ALIX antibodies (a gift from Dr. Wesley Sundquist, University of Utah) and biotinylated proteins were detected with streptavidin Alexa-Fluor-488 (Invitrogen S32354). Primary antibodies were detected with goat polyclonal Alexa Fluor-conjugated secondary antibodies (Invitrogen). Primary and secondary antibodies were both diluted in 3% BSA in PBS. Coverslips were incubated with primary antibodies for 1 hour at RT, washed, and incubated with secondary antibodies for 1 hour at RT. Vectashield with DAPI stain (Vector Laboratories) was used to mount the coverslips on slides. Fluorescence was detected using wide-field epifluorescence microscopy and images were analyzed using ImageJ.

### Gels and western blots

Cell lysates were prepared at the indicated time points post-infection in Laemmli sample buffer (BioRad). The samples were boiled for 5 min, separated on a Bolt 4-12% Bis-Tris gel (Invitrogen), and transferred to polyvinylidene difluoride (PVDF) membranes. Membranes were blocked with 5% nonfat dry milk in TBS supplemented with 0.5% Tween-20, and proteins were detected by incubation with primary antibodies diluted in blocking buffer followed by incubation with secondary antibodies (raised in goat against the appropriate species) conjugated to horseradish peroxidase (HRP) and diluted in blocking buffer. HA was detected using an HRP-conjugated HA antibody (Roche 12013819001), V5 was detected with mouse anti-V5 tag monoclonal antibody (Invitrogen R960), SAG2A was detected using rabbit polyclonal anti-SAG2A antibodies (46), MAF1b was detected using rabbit polyclonal anti-MAF1b antibodies (27), GAPDH was detected using mouse monoclonal anti-GAPDH antibody 6C5 (Calbiochem), and biotinylated proteins were detected using streptavidin-HRP (Invitrogen S911). HRP was detected using enhanced chemiluminescence (ECL) kit (Pierce). Silver-stained gels were generated using Pierce Silver Stain Kit (Thermo Scientific).

### Protein biotinylation

Biotin labeling in HFFs was initiated with the addition of 50 μM biotin (Sigma B4501; from a 100mM stock dissolved in DMSO) for 1 hour and terminated by cooling cells to 0°C on wet ice and washing away excess biotin (3-5 washes with ice cold PBS). The cells were then either fixed for subsequent staining for IFA, lysates prepared for western blot analysis (resuspended in cold RIPA and clarified with a 10,000 x g spin for 10 min at 4°C), or clarified lysates subjected to enrichment of the biotinylated proteins by streptavidin bead enrichment. To enrich for biotinylated proteins, the lysates were incubated with pre-washed Pierce streptavidin-coated magnetic beads (Thermo Fisher 88817) rotating overnight at 4°C. The beads were subsequently washed 2X with 1 mL of RIPA lysis buffer, 1X with 1 mL of 1 M KCl, 1X with 1 mL of 0.1 M Na2CO3, 1X with 1 mL of 2 M urea in 10 mM Tris-HCl (pH 8.0), and 3X with 1 mL RIPA lysis buffer. With the exception of the samples submitted for LC-MS/MS analysis, for which a detailed protocol is included below, the biotinylated proteins were eluted from the beads by incubation with Laemmli sample buffer supplemented with BME (BioRad) and 2 mM biotin at 90°C for 10 min.

### Sample preparation for proteomics

8 × 10^6^ HFFs were plated in eleven 15 cm dishes. 24 hours later, the HFFs were either infected with 12 x 10^6^ RHΔ*rop17*::ROP17-3xHA tachyzoites or mock-infected with an equivalent volume of syringe-lysed uninfected HFFs. 22 hours post-infection, the medium was replaced with either medium containing 50 μM biotin for 1 hour to initiate biotin labeling or with fresh medium containing no added biotin. After 1 hour of labeling, the cells were washed on wet ice 4X with 20 mL of ice cold PBS. 1 mL of RIPA lysis buffer supplemented with complete protease inhibitor cocktail (cOmplete, EDTA-free [Roche] and phosphatase inhibitor cocktail (PhosSTOP [Roche]) was added to each dish and the cells scraped into the lysis buffer and transferred into microfuge tubes. The lysates were then syringe-lysed 3X with a 27 G needle and subsequently clarified with a 10,000 x g spin for 10 min at 4°C. The supernatant was collected and total protein concentration was measured using the Pierce BCA Protein Assay Kit (Thermo Scientific).

For enrichment of biotinylated material, 3500 μg of each protein lysate was added to 250 μL of streptavidin-coated magnetic beads (pre-washed 5X with RIPA buffer) and allowed to incubate with rotation overnight at 4°C. The beads were subsequently washed 3X with 1 mL of RIPA lysis buffer (10 min each), 1X with 1 mL of 1 M KCl (10 min), 1X with 1 mL of 0.1 M Na_2_CO_3_ (1 min), 1X with 1 mL of 2 M urea in 10 mM Tris-HCl pH 8.0 (1 min), and 3X with 1 mL RIPA lysis buffer (10 min). The beads were transferred to a fresh microfuge tube prior to the final wash in RIPA buffer. 5% of the beads were removed and saved for quality control analysis of the enrichment. The protein from the 5% of beads was eluted by boiling the beads in Laemmli sample buffer supplemented with BME (BioRad) and 2 mM biotin at 90°C for 10 min. The remaining beads in RIPA lysis buffer were shipped overnight on ice to the Carr Laboratory (Broad Institute) for subsequent analysis by LC-MS/MS.

### On-bead trypsin digestion of biotinylated peptides

On-bead trypsin digestion was performed as described previously (18). The biotin-labeled proteins bound to magnetic beads were further washed to remove detergent traces. Magnetic beads were immobilized and washed twice with 200 μL of 50 mM Tris HCl buffer (pH 7.5) followed by two washes with 2 M urea/50 mM Tris (pH 7.5) buffer. A partial trypsin digestion was performed to release proteins from the beads by using 80 μL of 2 M urea/50 mM Tris containing 1 mM DTT and 0.4 μg trypsin (Mass Spectrometry Grade, Promega) for 1 h at 25 C. Magnetic beads were immobilized and the supernatant containing partially digested proteins was transferred to a fresh tube. The beads were washed twice with 60 μL of 2 M urea/50 mM Tris buffer (pH 7.5), and the washes were combined with the on-bead digest supernatant. Proteins were reduced with 4 mM DTT for 30 min at 25°C with shaking, followed by alkylation with 10 mM iodoacetamide for 45 min in the dark at 25°C. Proteins were completely digested by adding 0.5 μg of trypsin and incubating overnight at 25°C with shaking. Following the overnight incubation, samples were acidified to 1% formic acid (FA) and desalted using stage tips containing 2x C18 discs (Empore) as described next. The stage tip column was conditioned with 1x 100 μL methanol, 1x 100 μL 50% acetonitrile (ACN) / 1% FA, and 2x 100 μL 0.1% FA washes. Acidified peptides were bound to the column, washed with 2x 100 μL 0.1% FA, and eluted with 50 μL 50%ACN/0.1%FA. Eluted peptides were dried to using a vacuum concentrator.

### TMT labeling and fractionation of peptides

Desalted peptides were labeled with 11-plex TMT reagents (Thermo Fisher Scientific) and fractionated as described previously (18). Dried peptides were reconstituted in 100 μL of 50 mM HEPES, labeled using 0.8 mg of TMT reagent in 41 μL of anhydrous acetonitrile for 1 h at room temperature. The TMT reactions were quenched with 8 μL of 5% hydroxylamine at room temperature for 15 minutes with shaking, and the labeled peptides were desalted on C18 stage tips as described above. Peptides were fractionated by basic reverse phase using styrenedivinylbenzene-reverse phase sulfonate (SDB-RPS, Empore) material in stage tip columns. Two-disc punches were packed in a tip and equilibrated with 50 μL methanol, 50 μL 50% CAN / 0.1% FA, and 2x 75 μL 0.1% FA washes. Peptides were reconstituted in 0.1% FA and loaded into the column, followed by conditioning with 25 μL of 20 mM ammonium formate (AF). Next, peptides were sequentially eluted into 6x 100 μL fractions with 20mM AF and varying concentrations of ACN: 5%, 10%, 15%, 20%, 30%, and 55%. The six peptide fractions were dried by vacuum centrifugation.

### Liquid chromatography and mass spectrometry

Desalted peptides were resuspended in 9 μL of 3% ACN/0.1% FA and analyzed by online nanoflow liquid chromatography tandem mass spectrometry (LC-MS/MS) using a Proxeon Easy-nLC 1200 coupled to a Q Exactive HF-X Hybrid Quadrupole-Orbitrap Mass Spectrometer (Thermo Fisher Scientific). Four μL of sample from each fraction was separated on a capillary column (360 x 75 μm, 50 °C) containing an integrated emitter tip packed to a length of approximately 25 cm with ReproSil-Pur C18-AQ 1.9 μm beads (Dr. Maisch GmbH). Chromatography was performed with a 110 min gradient of solvent A (3% ACN/0.1% FA) and solvent B (90% ACN/0.1% FA). The gradient profile, described as min:% solvent B, was 0:2, 1:6, 85:30, 94:60, 95:90, 100:90, 101:50, 110:50. Ion acquisition was performed in data-dependent MS2 (ddMS2) mode with the following relevant parameters: MS1 acquisition (60,000 resolution, 3E6 AGC target, 10ms max injection time) and MS2 acquisition (Loop count = 20, 0.7m/z isolation window, 31 NCE, 45,000 resolution, 5E4 AGC target, 105 ms max injection time, 1E4 intensity threshold, 15s dynamic exclusion, and charge exclusion for unassigned, 1 and >6).

### Protein quantification

Collected RAW LC-MS/MS data were analyzed using Spectrum Mill software package v6.1 pre-release (Agilent Technologies). MS2 spectra were extracted from RAW files and merged if originating from the same precursor, or within a retention time window of +/− 60 s and m/z range of +/− 1.4, followed by filtering for precursor mass range of 750-6000 Da and sequence tag length > 0. MS/MS search was performed against a custom concatenated FASTA database containing (1) the human UniProt protein database downloaded December 2017, (2) a list of known common contaminants, and (3) the *Toxoplasma gondii* protein database downloaded from ToxoDB.org (version 43). Search parameters were set to “Trypsin allow P”, <5 missed cleavages, fixed modifications (cysteine carbamidomethylation and TMT11 on N-term and internal lysine), and variable modifications (oxidized methionine, acetylation of the protein N-terminus, pyroglutamic acid on N-term Q, and pyro carbamidomethyl on N-term C). Matching criteria included a 30% minimum matched peak intensity and a precursor and product mass tolerance of +/− 20 ppm. Peptide-level matches were validated if found to be below the 1.0% false discovery rate (FDR) threshold and within a precursor charge range of 2-6.

A protein-centric summary containing TMT channel intensities was exported from Spectrum Mill for downstream analysis in the R environment for statistical computing. Proteins were filtered to remove those that do not originate from humans or *Toxoplasma* and those with less than 2 unique peptides identified. Missing TMT intensity values were imputed assuming left-censored missing data by drawing values from the 1st percentile of all intensities. TMT ratios were calculated using the median of all TMT channels, followed by log_2_ transformation. Two-sample moderated t-tests were performed using the limma package with correction for multiple testing by calculating local FDR and q-values (47). The source code for the proteomics data analysis is available via a Github repository (https://github.com/pierremj/toxoplasma-proteomics).

### ROC threshold analysis

Receiver-operating characteristic (ROC) analysis was performed in R using the *rocit* and *plotROC* libraries. A list of known PV/PVM/exported *Toxoplasma* proteins were used as likely positives, while a list of predicted non-secreted *Toxoplasma* proteins [i.e. annotated as 19S/20S proteosome, 40S/60S ribosome, apicoplast, cytosolic, ER, mitochondrial, nuclear, plasma membrane, inner membrane complex, or tubulin cytoskeleton localized from a recent *Toxoplasma* LOPIT proteomics dataset (25), and not containing a predicted signal peptide, and not annotated as “Hypothetical”), were used as likely negatives. Both lists can be found in **File S1**. The log_2_ fold-change values ([mT-RAH + *Toxo*/mT +*Toxo*] and [mT-RAH +*Toxo*/mT-RAH *+Toxo,* no biotin]) were used as classifiers, and the optimal threshold points were determined using the Youden index method.

### Data analysis to generate tables

To generate **Table 1**, the *Toxoplasma* proteins from **File S1** were filtered based on: protein score >25, protein coverage >1%, mT-RAH/mT log_2_ FC >-0.65 (from ROC analysis), and mT-RAH +*Toxo*/mT-RAH *+Toxo* (no biotin) log_2_ FC >2.95 (from ROC analysis). All the *Toxoplasma* proteins that met these criteria are displayed in **Table 1**.

To generate **Table 2**, the human proteins from **File S1** were filtered based on: protein score >25, protein coverage >1%, mT-RAH/mT log_2_ FC >-0.65 (from ROC analysis), and mT-RAH +*Toxo*/mT-RAH *+Toxo* (no biotin) log_2_ FC >0 and adjusted p-value <0.01 (to remove endogenously biotinylated proteins). The list of proteins was then rank-sorted by the log_2_ FC of the mT-RAH + *Toxo*/mT-RAH +mock comparison, with the top 50 displayed in **Table 2**.

### Functional enrichment analysis

Functional enrichment analysis was performed using the *gost* function with default parameters from the *gprofiler2* library in R (48).

### STRING network analysis

Generation of the protein-protein interaction network was performed using StringApp (49) in Cytoscape (v. 3.8.2) (50). The network was filtered to remove singletons (proteins with no association to another protein) and to only include interactions for which experimental evidence existed within STRING (v. 11) (51) with high confidence (score >0.7).

## Data availability

Raw mass spectrometry data and processed Spectrum Mill files will be made publicly available in MassIVE upon acceptance of the manuscript.

## Acknowledgements

We thank all members of our laboratory for helpful discussions, especially Alma Mendoza for assistance with strain generation, Terence Theisen for help with data retrieval, and Melanie Espiritu for help with tissue culture and ordering. We also thank Wesley Sundquist for providing us with the anti-ALIX antibody, Jonathan Diep and Jan Carette for providing expertise and reagents for the generation of lentivirus, and the Carruthers, Weiss, Coppens, and Hanson labs for helpful discussions and exchange of information regarding host ESCRT.

This project has been funded in whole or part with federal funds from the National Institutes of Health under awards NIH R01-AI021423 (JCB), NIH R01-AI129529 (JCB), NIH R01-DK12140903 (AYT), NIH T32-AI732832 (AMC), NIH F32-

HL154711 (PMJB), and with funds from the National Science Foundation Graduate Research Fellowship Program (https://www.nsfgrfp.org/) under grant DGE-114747 (AMC). TCB was supported by Dow Graduate Research and Lester Wolfe Fellowships. The funders had no role in study design, data collection and analysis, or preparation of the manuscript.

## Author Contributions

AMC and JCB conceived the research and AMC performed all experiments except as noted in the following. AMC, JCB, PMJB, SAC, and TCB designed the proteomics experiment. AMC prepared the proteomics samples. PMJB processed the proteomic samples and performed mass spectrometry and quantitative data analysis. TCB and AYT shared technical expertise and unpublished reagents (plasmids). AMC drafted the manuscript and all authors reviewed and edited the manuscript.

## Figures and Figure Legends

**Supplemental File 1.** Complete quantitative mass spectrometry data.

**Supplemental File 2.** DNA sequences of fusion proteins used in this study.

**Figure S1.**
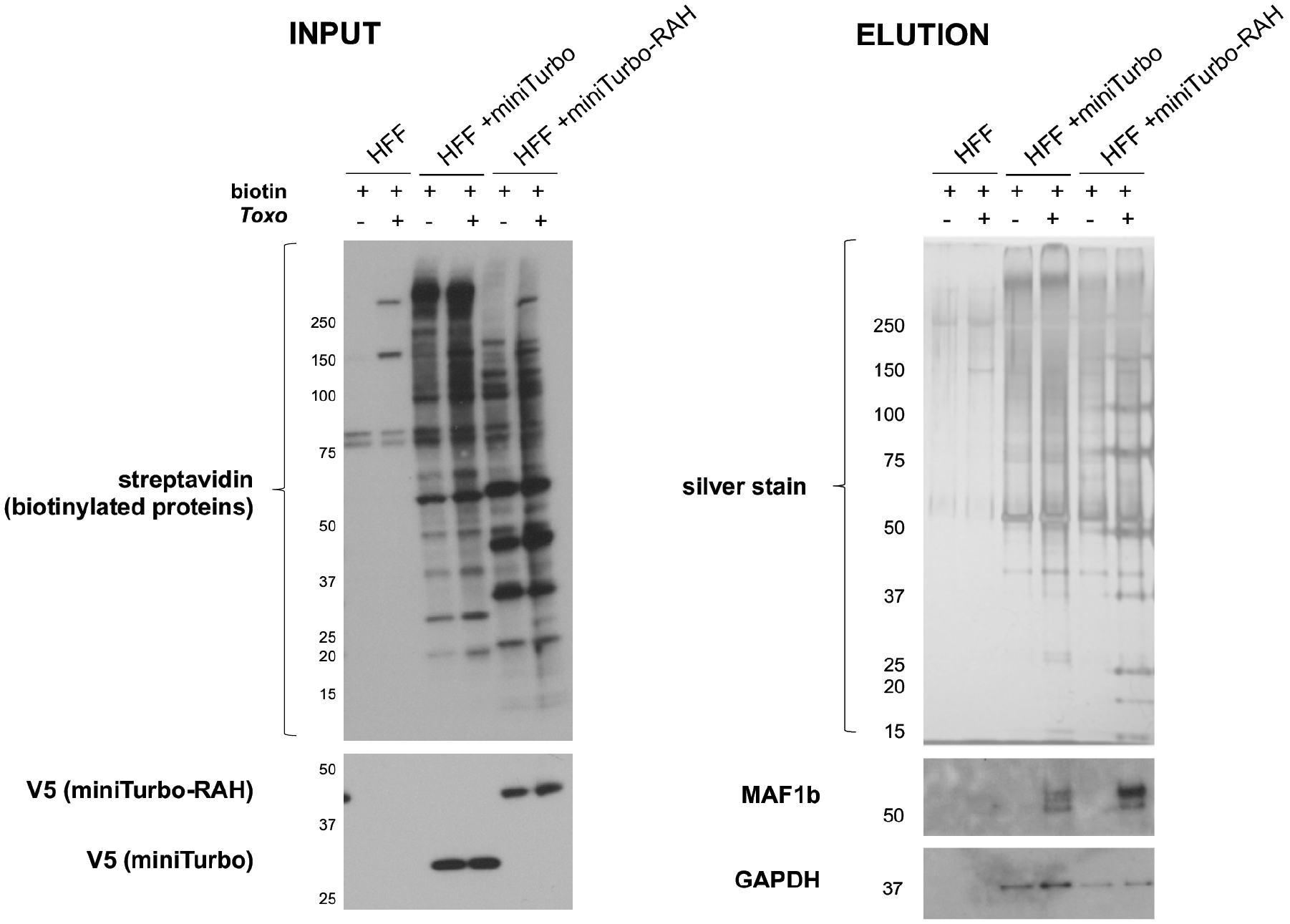
The *Toxoplasma* PVM-localized protein MAF1b is enriched in miniTurbo-RAH cells. INPUT: The indicated line of HFFs were either infected or left uninfected (+/−) with *RHΔku80Δhpt* tachyzoites for 24 hours and then incubated with or without (+/−) 50μg of biotin for 1 hour. Whole cell lysates were subsequently prepared and a sample run on SDS-PAGE gel and blotted with streptavidin-HRP to visualize all biotinylated proteins and mouse anti-V5 antibodies to visualize miniTurbo expression. ELUTION: Silver stain and western blots of eluates after streptavidin affinity purification. 100μg of each input lysate was incubated with streptavidin-coated magnetic beads overnight. The beads were washed and proteins eluted from the beads by boiling in the presence of 2mM biotin and BME. Half of each elution was run on an SDS-PAGE gel and blotted with rabbit anti-MAF1b antibodies and mouse anti-GAPDH antibodies to visualize enrichment of these proteins. The remaining halves of each elution were run on a separate SDS-PAGE gel and stained with silver to visualize total protein material eluted from the beads. Approximate migration of a ladder of size standards (kDa) is indicated.

**Figure S2.**
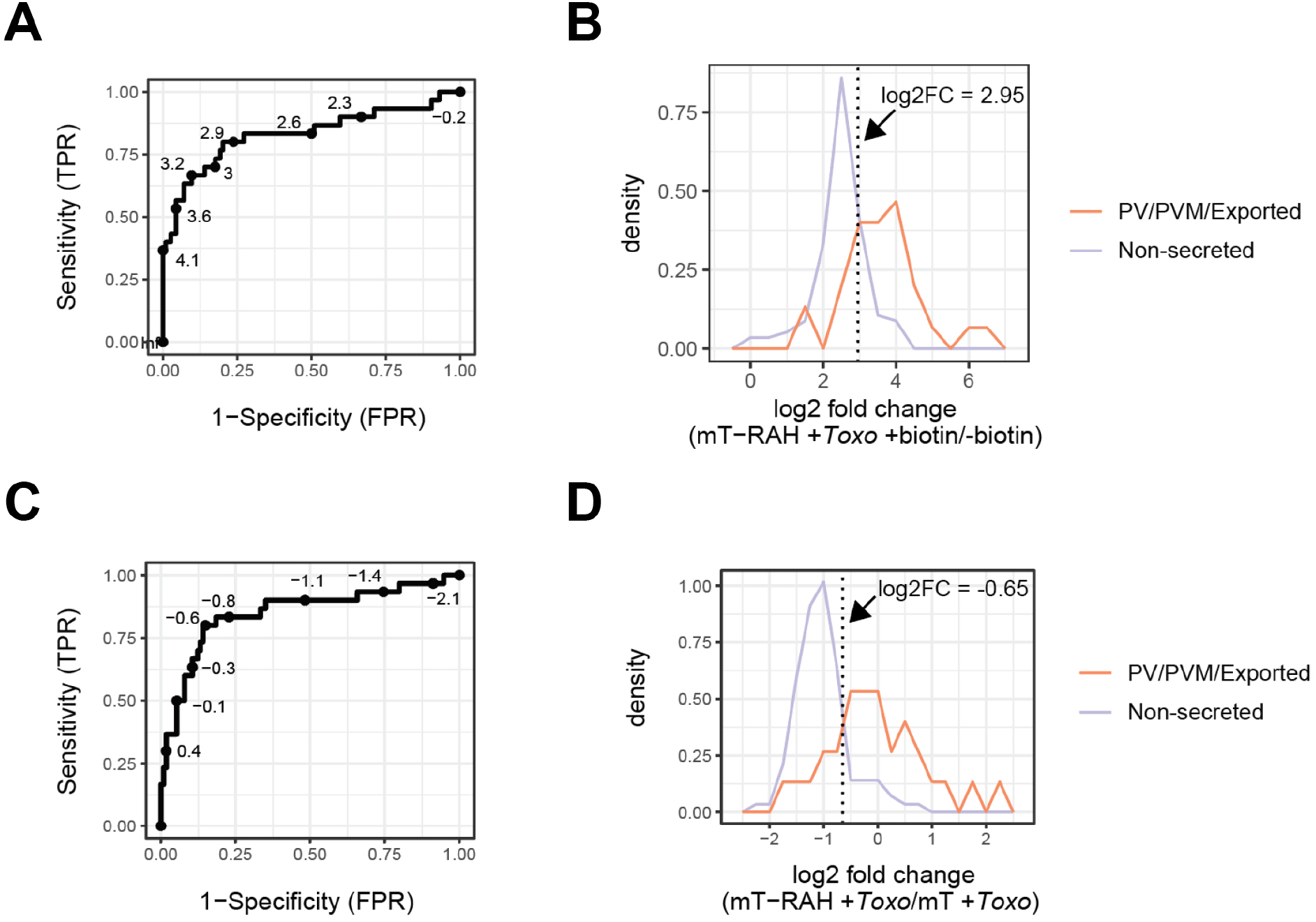
Analyses to determine optimal cutoff thresholds based on likely positives and likely negatives. **A.** Receiver operating characteristic (ROC) curve showing the true positive rate (TPR) and false positive rate (FPR) for classification of known PV/PVM/Exported-localized and putatively non-secreted *Toxoplasma* proteins (see Materials & Methods) at all log_2_ fold-change values ([mT-RAH + *Toxo*]/[mT-RAH *+ Toxo,* no biotin])). The optimal point (maximum value of Youden’s index, defined as max(J = sensitivity + specificity −1)) was identified as 2.95. **B.** Normalized density plot showing the frequency of known PV/PVM-localized and putatively non-secreted proteins across the log_2_ fold-change values ([mT-RAH + *Toxo*]/[mT-RAH *+ Toxo,* no biotin]). Dotted line indicates the optimal threshold as identified by ROC analysis in panel A. **C.** ROC curve generated as in panel A at all log_2_ fold-change values [mT-RAH +*Toxo*]/[mT +*Toxo*]. The optimal point (Youden’s index) was identified as −0.65. **D.** Normalized density plot showing the frequency of known PV/PVM-localized and putatively non-secreted proteins across the log_2_ fold-change values ([mT-RAH +*Toxo*]/[mT +*Toxo*]). Dotted line indicates the optimal threshold as identified by ROC analysis in panel C.

## References

1. Hill DE, Chirukandoth S, Dubey JP. 2005. Biology and epidemiology of Toxoplasma gondii in man and animals. Anim Heal Res Rev.

2. Melo EJ, Carvalho TM, De Souza W. 2001. Behaviour of microtubules in cells infected with Toxoplasma gondii. Biocell 25:53–59.

3. Romano JD, Bano N, Coppens I. 2008. New host nuclear functions are not required for the modifications of the parasitophorous vacuole of Toxoplasma. Cell Microbiol2007/11/01. 10:465–476.

4. Walker ME, Hjort EE, Smith SS, Tripathi A, Hornick JE, Hinchcliffe EH, Archer W, Hager KM. 2008. Toxoplasma gondii actively remodels the microtubule network in host cells. Microbes Infect2008/11/06. 10:1440–1449.

5. Jones TC, Hirsch JG. 1972. The interaction between Toxoplasma gondii and mammalian cells. II. The absence of lysosomal fusion with phagocytic vacuoles containing living parasites. J Exp Med 136:1173–1194.

6. Sinai AP, Webster P, Joiner KA. 1997. Association of host cell mitochondria and endoplasmic reticulum with the Toxoplasma gondii parasitophorous vacuole. J Cell Sci 110:2117–2128.

7. Pfefferkorn ER. 1984. Interferon gamma blocks the growth of Toxoplasma gondii in human fibroblasts by inducing the host cells to degrade tryptophan. Proc Natl Acad Sci U S A 81:908–912.

8. Coppens I, Sinai AP, Joiner KA. 2000. Toxoplasma gondii exploits host low-density lipoprotein receptor-mediated endocytosis for cholesterol acquisition. J Cell Biol 149:167–180.

9. Dimier IH, Bout DT. 1998. Interferon-gamma-activated primary enterocytes inhibit Toxoplasma gondii replication: a role for intracellular iron. Immunology 94:488–495.

10. Hakimi MA, Olias P, Sibley LD. 2017. Toxoplasma effectors targeting host signaling and transcription. Clin Microbiol Rev.

11. Panas MW, Boothroyd JC. 2020. Toxoplasma Uses GRA16 To Upregulate Host c-Myc. mSphere.

12. Etheridge RD, Alaganan A, Tang K, Lou HJ, Turk BE, Sibley LD. 2014. The Toxoplasma pseudokinase ROP5 forms complexes with ROP18 and ROP17 kinases that synergize to control acute virulence in mice. Cell Host Microbe 15:537–550.

13. Kim EW, Nadipuram SM, Tetlow AL, Barshop WD, Liu PT, Wohlschlegel JA, Bradley PJ. 2016. The Rhoptry Pseudokinase ROP54 Modulates Toxoplasma gondii Virulence and Host GBP2 Loading. mSphere 1.

14. Rastogi S, Cygan AM, Boothroyd JC. 2019. Translocation of effector proteins into host cells by Toxoplasma gondii. Curr Opin Microbiol.

15. Clough B, Frickel EM. 2017. The Toxoplasma Parasitophorous Vacuole: An Evolving Host-Parasite Frontier. Trends Parasitol.

16. Nadipuram SM, Kim EW, Vashisht AA, Lin AH, Bell HN, Coppens I, Wohlschlegel JA, Bradley PJ. 2016. In Vivo biotinylation of the toxoplasma parasitophorous vacuole reveals novel dense granule proteins important for parasite growth and pathogenesis. MBio 7.

17. Pan M, Li M, Li L, Song Y, Hou L, Zhao J, Shen B. 2018. Identification of Novel Dense-Granule Proteins in Toxoplasma gondii by Two Proximity-Based Biotinylation Approaches. J Proteome Res.

18. Branon TC, Bosch JA, Sanchez AD, Udeshi ND, Svinkina T, Carr SA, Feldman JL, Perrimon N, Ting AY. 2018. Efficient proximity labeling in living cells and organisms with TurboID. Nat Biotechnol.

19. Roux KJ, Kim DI, Raida M, Burke B. 2012. A promiscuous biotin ligase fusion protein identifies proximal and interacting proteins in mammalian cells. J Cell Biol 196:801–810.

20. Reese ML, Boothroyd JC. 2009. A helical membrane-binding domain targets the Toxoplasma ROP2 family to the parasitophorous vacuole. Traffic 10:1458–1470.

21. Sibley LD, Niesman IR, Parmley SF, Cesbron-Delauw MF. 1995. Regulated secretion of multi-lamellar vesicles leads to formation of a tubulovesicular network in host-cell vacuoles occupied by Toxoplasma gondii. J Cell Sci 108:1669–1677.

22. Fricker M, Hollinshead M, White N, Vaux D. 1997. Interphase nuclei of many mammalian cell types contain deep, dynamic, tubular membrane-bound invaginations of the nuclear envelope. J Cell Biol.

23. Jelenska J, Crawford MJ, Harb OS, Zuther E, Haselkorn R, Roos DS, Gornicki P. 2001. Subcellular localization of acetyl-CoA carboxylase in the apicomplexan parasite Toxoplasma gondii. Proc Natl Acad Sci U S A 98:2723–2728.

24. Drewry LL, Jones NG, Wang Q, Onken MD, Miller MJ, Sibley LD. 2019. The secreted kinase ROP17 promotes Toxoplasma gondii dissemination by hijacking monocyte tissue migration. Nat Microbiol.

25. Barylyuk K, Koreny L, Ke H, Butterworth S, Crook OM, Lassadi I, Gupta V, Tromer E, Mourier T, Stevens TJ, Breckels LM, Pain A, Lilley KS, Waller RF. 2020. A Comprehensive Subcellular Atlas of the Toxoplasma Proteome via hyperLOPIT Provides Spatial Context for Protein Functions. Cell Host Microbe.

26. Hakansson S, Charron AJ, Sibley LD. 2001. Toxoplasma evacuoles: a two-step process of secretion and fusion forms the parasitophorous vacuole. Embo J 20:3132–3144.

27. Pernas L, Adomako-Ankomah Y, Shastri AJ, Ewald SE, Treeck M, Boyle JP, Boothroyd JC. 2014. Toxoplasma Effector MAF1 Mediates Recruitment of Host Mitochondria and Impacts the Host Response. PLoS Biol 12.

28. Lecordier L, Mercier C, Torpier G, Tourvieille B, Darcy F, Liu JL, Maes P, Tartar A, Capron A, Cesbron-Delauw MF. 1993. Molecular structure of a Toxoplasma gondii dense granule antigen (GRA 5) associated with the parasitophorous vacuole membrane. Mol Biochem Parasitol 59:143–153.

29. El Hajj H, Lebrun M, Fourmaux MN, Vial H, Dubremetz JF. 2007. Inverted topology of the Toxoplasma gondii ROP5 rhoptry protein provides new insights into the association of the ROP2 protein family with the parasitophorous vacuole membrane. Cell Microbiol 9:54–64.

30. Labruyere E, Lingnau M, Mercier C, Sibley LD. 1999. Differential membrane targeting of the secretory proteins GRA4 and GRA6 within the parasitophorous vacuole formed by Toxoplasma gondii. Mol Biochem Parasitol 102:311–324.

31. Carey KL, Donahue CG, Ward GE. 2000. Identification and molecular characterization of GRA8, a novel, proline-rich, dense granule protein of Toxoplasma gondii. Mol Biochem Parasitol 105:25–37.

32. Tu V, Ma Y, Tomita T, Sugi T, Mayoral J, Han B, Yakubu RR, Williams T, Horta A, Weiss LM. 2020. The Toxoplasma gondii cyst wall interactome. MBio.

33. Käll L, Krogh A, Sonnhammer ELL. 2007. Advantages of combined transmembrane topology and signal peptide prediction-the Phobius web server. Nucleic Acids Res.

34. Wang Y, Sangaré LO, Paredes-Santos TC, Krishnamurthy S, Hassan MA, Furuta AM, Markus BM, Lourido S, Saeij JPJ. 2019. A genome-wide loss-of-function screen identifies *Toxoplasma gondii* genes that determine fitness in interferon gamma-activated murine macrophages. bioRxiv 867705.

35. Vietri M, Radulovic M, Stenmark H. 2020. The many functions of ESCRTs. Nat Rev Mol Cell Biol.

36. Guérin A, Corrales RM, Parker ML, Lamarque MH, Jacot D, El Hajj H, Soldati-Favre D, Boulanger MJ, Lebrun M. 2017. Efficient invasion by Toxoplasma depends on the subversion of host protein networks. Nat Microbiol 2:1358–1366.

37. Maki M, Takahara T, Shibata H. 2016. Multifaceted roles of ALG-2 in Ca2+-regulated membrane trafficking. Int J Mol Sci.

38. Di Mattia T, Wilhelm LP, Ikhlef S, Wendling C, Spehner D, Nominé Y, Giordano F, Mathelin C, Drin G, Tomasetto C, Alpy F. 2018. Identification of MOSPD2, a novel scaffold for endoplasmic reticulum membrane contact sites. EMBO Rep.

39. Christ L, Raiborg C, Wenzel EM, Campsteijn C, Stenmark H. 2017. Cellular Functions and Molecular Mechanisms of the ESCRT Membrane-Scission Machinery. Trends Biochem Sci.

40. Martinelli N, Hartlieb B, Usami Y, Sabin C, Dordor A, Miguet N, Avilov S V., Ribeiro EA, Göttlinger H, Weissenhorn W. 2012. CC2D1A is a regulator of ESCRT-III CHMP4B. J Mol Biol.

41. Yin B, Mendez R, Zhao XY, Rakhit R, Hsu KL, Ewald SE. 2020. Automated Spatially Targeted Optical Microproteomics (autoSTOMP) to Determine Protein Complexity of Subcellular Structures. Anal Chem.

42. Panas MW, Ferrel A, Naor A, Tenborg E, Lorenzi HA, Boothroyd JC. 2019. Translocation of Dense Granule Effectors across the Parasitophorous Vacuole Membrane in Toxoplasma-Infected Cells Requires the Activity of ROP17, a Rhoptry Protein Kinase. mSphere.

43. Fox BA, Ristuccia JG, Gigley JP, Bzik DJ. 2009. Efficient gene replacements in Toxoplasma gondii strains deficient for nonhomologous end joining. Eukaryot Cell2009/02/17. 8:520–529.

44. Marino ND, Panas MW, Franco M, Theisen TC, Naor A, Rastogi S, Buchholz KR, Lorenzi HA, Boothroyd JC. 2018. Identification of a novel protein complex essential for effector translocation across the parasitophorous vacuole membrane of Toxoplasma gondii. PLoS Pathog 14:e1006828.

45. Saeij JPJ, Boyle JP, Coller S, Taylor S, Sibley LD, Brooke-Powell ET, Ajioka JW, Boothroyd JC. 2006. Polymorphic Secreted Kinases Are Key Virulence Factors in Toxoplasmosis. Science (80−) 314:1780–1783.

46. Grigg ME, Bonnefoy S, Hehl AB, Suzuki Y, Boothroyd JC. 2001. Success and virulence in Toxoplasma as the result of sexual recombination between two distinct ancestries. Science (80−) 294:161–165.

47. Storey JD. 2002. A direct approach to false discovery rates. J R Stat Soc Ser B Stat Methodol.

48. Raudvere U, Kolberg L, Kuzmin I, Arak T, Adler P, Peterson H, Vilo J. 2019. G:Profiler: A web server for functional enrichment analysis and conversions of gene lists (2019 update). Nucleic Acids Res.

49. Doncheva NT, Morris JH, Gorodkin J, Jensen LJ. 2019. Cytoscape StringApp: Network Analysis and Visualization of Proteomics Data. J Proteome Res.

50. Shannon P, Markiel A, Ozier O, Baliga NS, Wang JT, Ramage D, Amin N, Schwikowski B, Ideker T. 2003. Cytoscape: A software Environment for integrated models of biomolecular interaction networks. Genome Res.

51. Szklarczyk D, Gable AL, Lyon D, Junge A, Wyder S, Huerta-Cepas J, Simonovic M, Doncheva NT, Morris JH, Bork P, Jensen LJ, Von Mering C. 2019. STRING v11: Protein-protein association networks with increased coverage, supporting functional discovery in genome-wide experimental datasets. Nucleic Acids Res.

